# *Aspergillus fumigatus* G-protein coupled receptors GprM and GprJ are important for the regulation of the cell wall integrity pathway, secondary metabolite production, and virulence

**DOI:** 10.1101/2020.08.03.235119

**Authors:** Aílton Pereira da Costa Filho, Guilherme Thomaz Pereira Brancini, Patrícia Alves de Castro, Jaire Alves Ferreira, Lilian Pereira Silva, Marina Campos Rocha, Iran Malavazi, João Guilherme de Moraes Pontes, Taícia Fill, Roberto Nascimento Silva, Fausto Almeida, Jacob L. Steenwyk, Antonis Rokas, Thaila F. dos Reis, Laure N.A. Ries, Gustavo H. Goldman

## Abstract

G-protein coupled receptors (GPCRs) are extracellular signalling receptors that sense environmental cues to coordinate a biological response. Fungi sense their environment primarily through GPCR-mediated signalling pathways, which in turn regulate fungal development, metabolism, virulence and mycotoxin biosynthesis. *A. fumigatus* is an important human pathogen that causes aspergillosis, a heterogeneous group of diseases that presents a wide range of clinical manifestations. Here, we investigate in detail the role of the GPCRs GprM and GprJ in growth and gene expression. GprM and GprJ are important for melanin production and the regulation of the cell wall integrity (CWI) pathway. Overexpression of *gprM* and *gprJ* causes a 20 and 50% reduction in growth rate when compared to the wild-type (WT) strain, and increases sensitivity to cell wall-damaging agents. Phosphorylation of the CWI protein kinase MpkA is increased in the Δ*gprM* and Δ*gprJ* strains and decreased in the overexpression mutants when compared to the WT strain. Furthermore, differences in cell wall polysaccharide concentrations and organization were observed in these strains. RNA-sequencing suggests that GprM and GprJ negatively regulate genes encoding secondary metabolites (SMs). Mass spectrometry analysis confirmed that the production of fumagillin, pyripyropene, fumigaclavine C, fumiquinazoline, and fumitremorgin is reduced in the Δ*gprM* and Δ*gprJ* strains, and that this regulation occurs, at least partially, through the activation of MpkA. Overexpression of *grpM* also resulted in the regulation of many transcription factors, with AsgA predicted to function downstream of GprM and MpkA signalling. Finally, we show that the Δ*gprM* and Δ*gprJ* mutants are reduced in virulence in the *Galleria mellonella* insect model of invasive aspergillosis. This work further contributes to unravelling functions of *A. fumigatus* GPCRs and shows that GprM and GprJ are essential for CWI, secondary metabolite production and virulence.

**Author summary:** *A. fumigatus* is the main ethiological agent of invasive pulmonary aspergillosis, a life-threatening fungal disease that occurs in severely immuno-compromised humans. Withstanding the host environment is essential for *A. fumigatus* virulence and sensing of extracellular cues occurs primarily through G-protein coupled receptors (GPCRs) that activate signal transduction pathways, which in turn regulate fungal development, metabolism, virulence and mycotoxin biosynthesis. The *A. fumigatus* genome encodes 15 putative classical GPCRs, with only three having been functionally characterized to date. In this work, we show that the two GPCRs GprM and GprJ regulate the phosphorylation of the mitogen-activated protein kinase MpkA and thus control the regulation of the cell wall integrity pathway. GprM and GprJ are also involved in the regulation of the production of the secondary metabolites fumagillin, pyripyropene, fumigaclavine C, fumiquinazoline, melanin, and fumitremorgin and this regulation partially occurs through the activation of MpkA. Furthermore, GprM and GprJ are important for virulence in the insect model *Galleria mellonella*. This work therefore functionally characterizes two GPCRs and shows how they regulate several intracellular pathways that have been shown to be crucial for *A. fumigatus* virulence.

## Introduction

*A. fumigatus* is an important human pathogen that causes aspergillosis, a heterogeneous group of diseases that presents a wide range of clinical manifestations, including chronic pulmonary aspergillosis (CPA), allergic bronchopulmonary aspergillosis (ABPA) and invasive pulmonary aspergillosis (IPA) [1]. IPA is the most serious pathology in terms of patient outcome and treatment. It primarily affects those who are severely immunocompromised such as patients with cancer, hematological neoplasms or those undergoing chemotherapy [1, 2] and is accompanied by a mortality rate ranging between 50% and 100% [3].

In eukaryotes, heterotrimeric G-protein-mediated signaling pathways play a pivotal role in transmembrane signaling. G-protein coupled receptors (GPCRs) are extracellular signalling receptors that sense environmental cues which initiate intracellular G-protein signalling, to coordinate a biological response. GPCRs are generally characterized by the presence of seven transmembrane (TM) domains and their association with intracellular G-proteins. The binding of an extracellular ligand to the receptor initiates intracellular signalling by stimulating the associated heterotrimeric G-proteins to exchange GDP for GTP, causing them to dissociate into the Gα subunit and a Gβ/Gγ dimer, which each have their own function in the activation or inactivation of specific pathways. G-protein functions are transient, and are regulated by repressors of G-protein signalling (RGS), which promote G-protein re-association, RGS ubiquitination and other feedback mechanisms [4].

Fungi sense their environment primarily through GPCR-mediated signalling pathways, which in turn regulate fungal development, metabolism, virulence and mycotoxin biosynthesis. Fungal GPCRs are able to detect hormones, proteins, nutrients, ions, hydrophobic surfaces and light [5]. Interestingly, fungal GPCRs do not belong to any of the mammalian receptor classes, making them specific targets for controlling fungal disease [6]. There are a lot of differences in the abundance and diversity of these receptors in fungi, and the potential ligands they detect. In the subkingdom of dikarya, the subphylum Pezizomycotina contains a higher number and diversity of classical and non-classical GPCRs than Saccharomycotina and Basidiomycota [for a review, see 7]. Among the *Aspergillus spp* Pezizomycotina, such as *A. nidulans, A. flavus*, and *A. fumigatus*, the number of classical GPCRS in each species is highly conserved [7].

In *A. fumigatus*, 161 proteins were found to encode seven predicted transmembrane domains (TMDs) [8]; however, 15 putative classical GPCRs have been identified in this pathogenic fungus [8, 9]. Deletion of *gprC* and *gprD* resulted in strains with severe growth defects as hyphal extension was reduced, germination was retarded, and hyphae showed elevated levels of branching [10]. Furthermore, the Δ*gprC* and Δ*gprD* strains were more sensitive to oxidative stress, more tolerant to cyclosporine (an inhibitor of the protein phosphatase calcineurin) and displayed attenuation of virulence in a murine infection model [10]. Deletion of *gprK* resulted in a strain with reduced conidiation and increased germination rate, and GprK was shown to control fungal development through the cAMP-PKA pathway to regulate the expression of developmental genes [11]. GprK was also proposed to be involved in sensing pentose sugars and in the control of the oxidative stress response, through regulating the expression of catalase- and superoxide dismutase-encoding genes via the mitogen-activated protein kinase (MAPK) SakA signalling pathway [11]. In addition, GprK is important for the production of the SM gliotoxin, although this GPCR was dispensable for virulence in the *G. mellonella* insect model of invasive aspergillosis [11].

Recently, we showed that the GPCR GprM is a negative regulator of melanin production as its deletion results in an increase in the melanin dihydroxynaphthalene (DHN) [12]. Melanins are a class of dark-brown pigments often associated with the cell wall. Their main role is to protect the organisms from exogenous stressors, thereby contributing to the first line of defense against external hazards [13, 14]. *A. fumigatus* produces two types of melanins: pyomelanin, which is derived from the catabolism of tyrosine via the intermediate homogentisate, and DHN-melanin, which is produced as a polyketide derivative and is responsible for the gray-green color of the spores [13]. Similar phenotypes were observed in strains deleted for the Gα protein GpaA and the MAPK MpkB [12], suggesting that these three proteins function in the same pathway. Split-ubiquitin based membrane yeast two-hybrid and co-immunoprecipitation assays confirmed that GpaA and GprM interact, suggesting their role in the MpkB signaling cascade [12].

Apart from the aforementioned studies, other *A. fumigatus* GPCRs have not been functionally characterized and we currently have very limited understanding on how this pathogenic fungus can sense host cues. In this work, we further investigated the role of GprM and also of GprJ in fungal growth, gene expression, cell wall integrity, SM production and virulence. Both GPCRs are important for the regulation of the cell wall integrity (CWI) pathway and the production of several SMs in addition to contributing to *A. fumigatus* virulence.

## Results

### The GPCRs GprM and GprJ are important for melanin production and the regulation of the CWI pathway

Previously, we have shown that the MAPK MpkB is important for conidiation and that its deletion increases dihydroxynaphthalene (DHN)-melanin production, as observed by a dark colouring of liquid culture supernatants [12]. Culture supernatants of strains deleted for the Gα protein GpaA and for the G protein-coupled receptor GprM also turned dark [12]. To identify additional pathways involved in DHN-melanin regulation, we exploited the dark-color production in the supernatant as a readout system. Using the same methodology, we screened 12 *A. fumigatus* GPCR deletion strains (Supplementary Table S1) in order to determine whether additional GPCRs are involved in melanin production. Dark-coloured supernatants were observed for the Δ*gprM* and *ΔgprJ* strains when compared to the wild-type (WT) strain but not for the Δ*gprM ΔgprJ* strain (Supplementary Figure S1). Complementation of Δ*gprJ* and simultaneous deletion of the *pksP* gene, encoding the polyketide synthase PksP required for DHN-melanin biosynthesis, confirm that the change in culture medium was linked to DHN-melanin (Supplementary Figure S1). *In silico* homology and similarity analyses, group GprM and GprJ as class 7 (orthologues of MG00532 from *Magnaporthe oryzae* with weak similarity to the rat growth-hormone-releasing factor) and class 4 (nitrogen) GPCRs, respectively [7] (Figure 1A). GprM and GprJ are present in 45 *Aspergillus* species (except *A. awamori* for GprJ) and both GPCRs are present in five different *A. fumigatus* strains (Figure 1B). In addition to the *gprM* and *gprJ* single and double deletion strains, we constructed the corresponding *gprM* and *gprJ* overexpression strains (Supplementary Table S1). These strains have comparable growth and conidiation to the wild-type strain on solid (Figure 1C) and in liquid (Supplementary Figure S2A) minimal medium (MM). The overexpression strains (Supplementary Table S1) were constructed by replacing the *gprM* and *gprJ* endogenous promoter by the inducible *xylp* promoter from *Penicillium chrysogenum*, which is induced by xylose and repressed by glucose [15]. Overexpression was confirmed by qRT-PCR (Figures 2A and 3A). Induction of *gprM* was increased 66- and 106-fold in the *xylp::gprM* strain when grown for 2 and 4 h in xylose, respectively, in comparison to the WT strain (Figure 2A). Induction of *gprJ* was increased 100- and 260-fold in the *xylp::gprJ* strain when grown for 30 min and 1 h in xylose-rich medium in comparison to the WT strain (Figure 3A).

**Figure 1.**
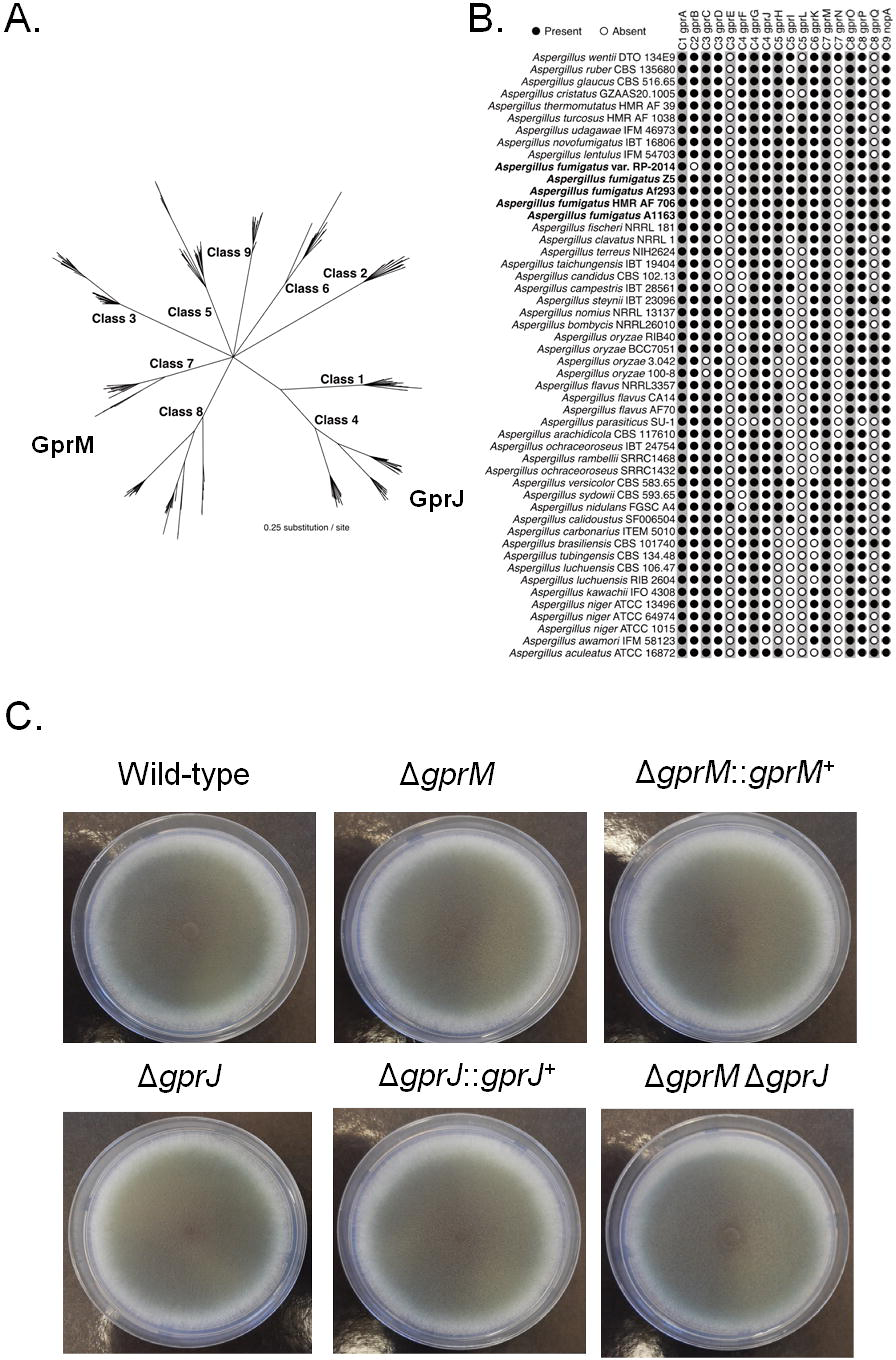
*A. fumigatus* GprM and GprJ homologues are present in different *Aspergillus spp.* A. Phylogenetic relationship among the nine classes of GPCRs present in *Aspergilli*. GprM is a class 7 GPCR whereas GprJ is a class 4 GPCR. B. Distribution of 18 *A. fumigatus* GPCR homologues in *Aspergillus* species. *A. fumigatus* strains are highlighted in bold. The presence of a GPCR is marked by a black circle whereas the absence is indicated by a white circle. C. The knock-out of *gprM* and *gprJ* in single and double deletion strains does not affect fungal growth on glucose minimal medium. Strains were grown from 10^5^ spores for 5 days at 37°C.

**Figure 2.**
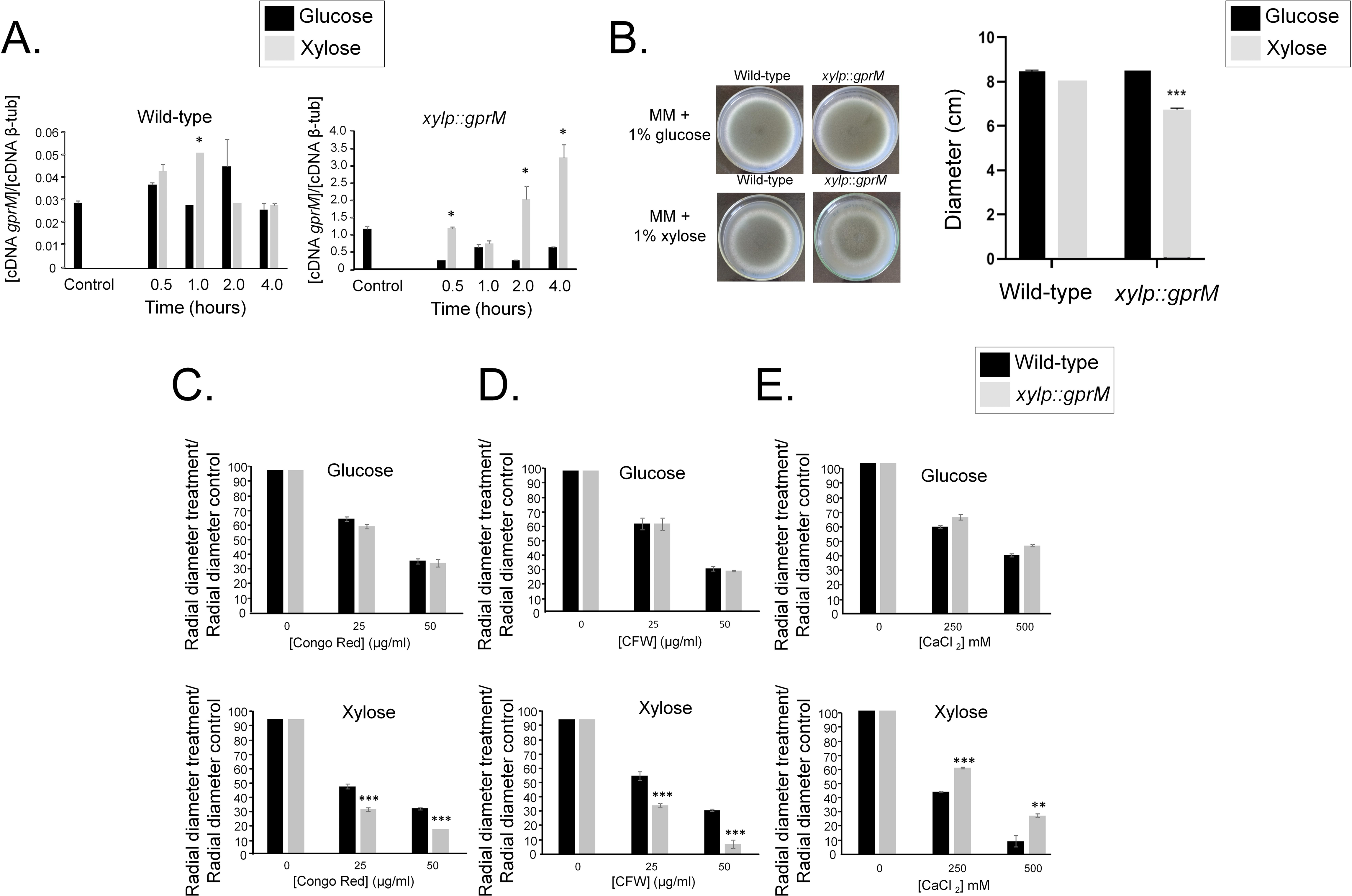
Overexpression of *gprM* causes decreased growth fitness and increased sensitivity to cell wall damaging agents. A. Overexpression of *gprM* was confirmed by RT-qPCR. Strains were grown for 24 h at 37°C in glucose minimal medium (GMM) (control) and transferred to xylose minimal medium (XMM) for 0.5, 1, 2 and 4 h before RNA was extracted and reverse transcribed to cDNA. Copy numbers of *gprM* were normalised by β-tubulin (Afu1g10910). B. Overexpression of *gprM* affects growth. Strains were grown from 10^5^ spores for 5 days at 37°C on GMM or XMM, before radial diameter was measured. Pictures of left panel are indicative for measurements of the graph shown in the right pannel. C. - E. Overexpression of *gprM* increases sensitivity to cell wall damagaing agents. Strains were grown from 10^5^ spores for 5 days at 37°C on GMM or XMM supplemented with increasing concentrations of congo red (C), calcofluor white (CFW, D) and calcium chloride (CaCl2) (E). Growth was normalised by growth in the control, drug-free condition. Standard deviations present the average of three independent biological repetitions with *p-value < 0.05, **p-value < 0.005 and ***p-value < 0.0005 in a one-way ANOVA with Tukey’s test for Post-Hoc analysis when compared to the wild-type strain.

**Figure 3.**
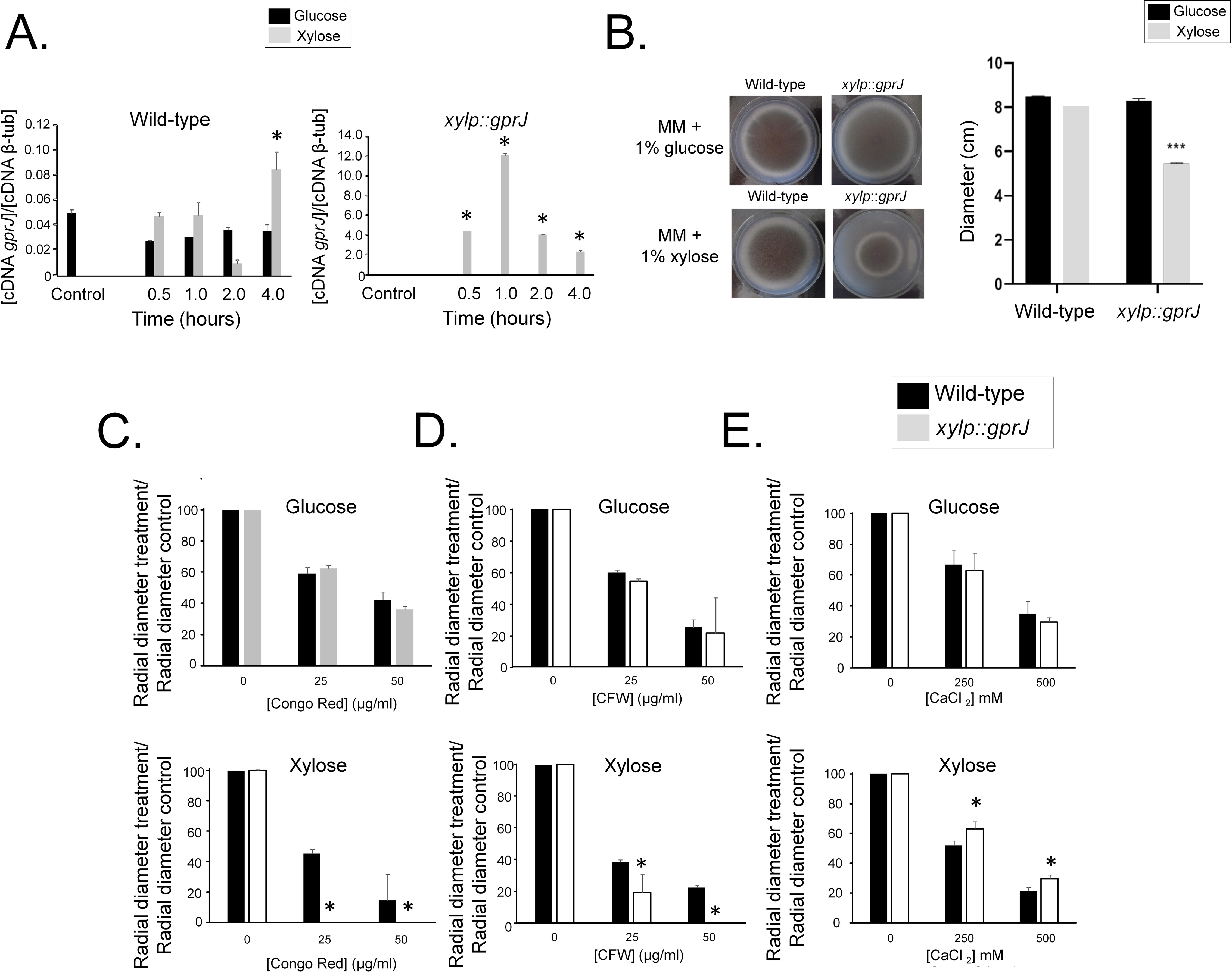
Overexpression of *gprJ* causes decreased growth fitness and increased sensitivity to cell wall damaging agents. A. Overexpression of *gprJ* was confirmed by RT-qPCR. Strains were grown for 24 h at 37°C in glucose minimal medium (GMM) (control) and transferred to xylose minimal medium (XMM) for 0.5, 1, 2 and 4 h before RNA was extracted and reverse transcribed to cDNA. Copy numbers of *gprM* were normalised by β-tubulin (Afu1g10910). B. Overexpression of *gprJ* affects growth. Strains were grown from 10^5^ spores for 5 days at 37°C on GMM or XMM, before radial diameter was measured. Pictures of left panel are indicative for measurements of the graph shown in the right pannel. (C–E) Overexpression of *gprJ* increases sensitivity to cell wall damaging agents. Strains were grown from 10^5^ spores for 5 days at 37°C on GMM or XMM supplemented with increasing concentrations of congo red (C), calcofluor white (CFW, D) and calcium chloride (CaCl_2_) (E). Growth was normalised by growth in the control, drug-free condition. Standard deviations present the average of three independent biological repetitions with *p-value < 0.05, **p-value < 0.005 and ***p-value < 0.0005 in a one-way ANOVA with Tukey’s test for Post-Hoc analysis when compared to the wild-type strain.

We then grew all strains in glucose minimal medium (GMM) supplemented with increasing concentrations of cell wall-perturbing agents. All growth was normalized in comparison to the drug-free, control condition. The Δ*gprM*, Δ*gprJ*, and Δ*gprM* Δ*gprJ* strains were not sensitive to these compounds. In contrast, the *gprM* overexpression strain has a 20 % reduction in growth when compared to the WT strain when grown on xylose minimal medium (XMM) (Figure 2B). Furthermore, *gprM* overexpression causes a significant reduction in growth in the presence of Congo Red (CR), and Calcofluor White (CFW) (Figures 2C and 2D). The *xylP*::*gprM* strain also presented increased growth in the presence of CaCl_2_ (Figure 2E). Similar to the *xylP::gprM* strain, overexpression of *gprJ* resulted in 50 % reduced growth when compared to the wild-type strain in the presence of XMM (Figure 3B); and in significantly reduced growth in the presence of CR and CFW (Figures 3C and 3D) and increased growth in the presence of CaCl_2_ (Figure 3E).

Due to the increased sensitivity of the *gprM* and *gprJ* overexpression strains to cell wall perturbing compounds, we further investigated the role of GprM and GprJ in cell wall maintenance. *A. fumigatus* MpkA is the main MAPK responsible for the CWI pathway and cell wall remodelling with MpkA phosphorylation signalling CWI pathway activation [16]. We investigated the impact of the deletion and overexpression of *gprM* and *gprJ* on MpkA phosphorylation (Figure 4). We used the Δ*mpkB* strain as a control because it was shown to have increased MpkA phosphorylation after 48 h growth in GMM when compared to the WT strain [12] (Figure 4A). To normalize the observed strain-specific phosphorylated MpkA levels, we used anti-ß-actin as the antibody that detects total cellular protein since the anti-p42-44 antibody to detect total MpkA does not function for *A. fumigatus* cellular protein extracts. MpkA phosphorylation was increased after 24 and 48 h growth in GMM in the Δ*gprM* and Δ*gprJ* strains when compared to the WT strain (Figure 4A). In contrast, overexpression of *gprM* and *gprJ* caused a decrease in MpkA phosphorylation when compared to the WT strain (Figure 4B). The Δ*gprM* Δ*gprJ* strain had reduced MpkA phosphorylation when compared to the WT strain (data not shown), suggesting that they genetically interact in the regulation of the CWI pathway.

**Figure 4.**
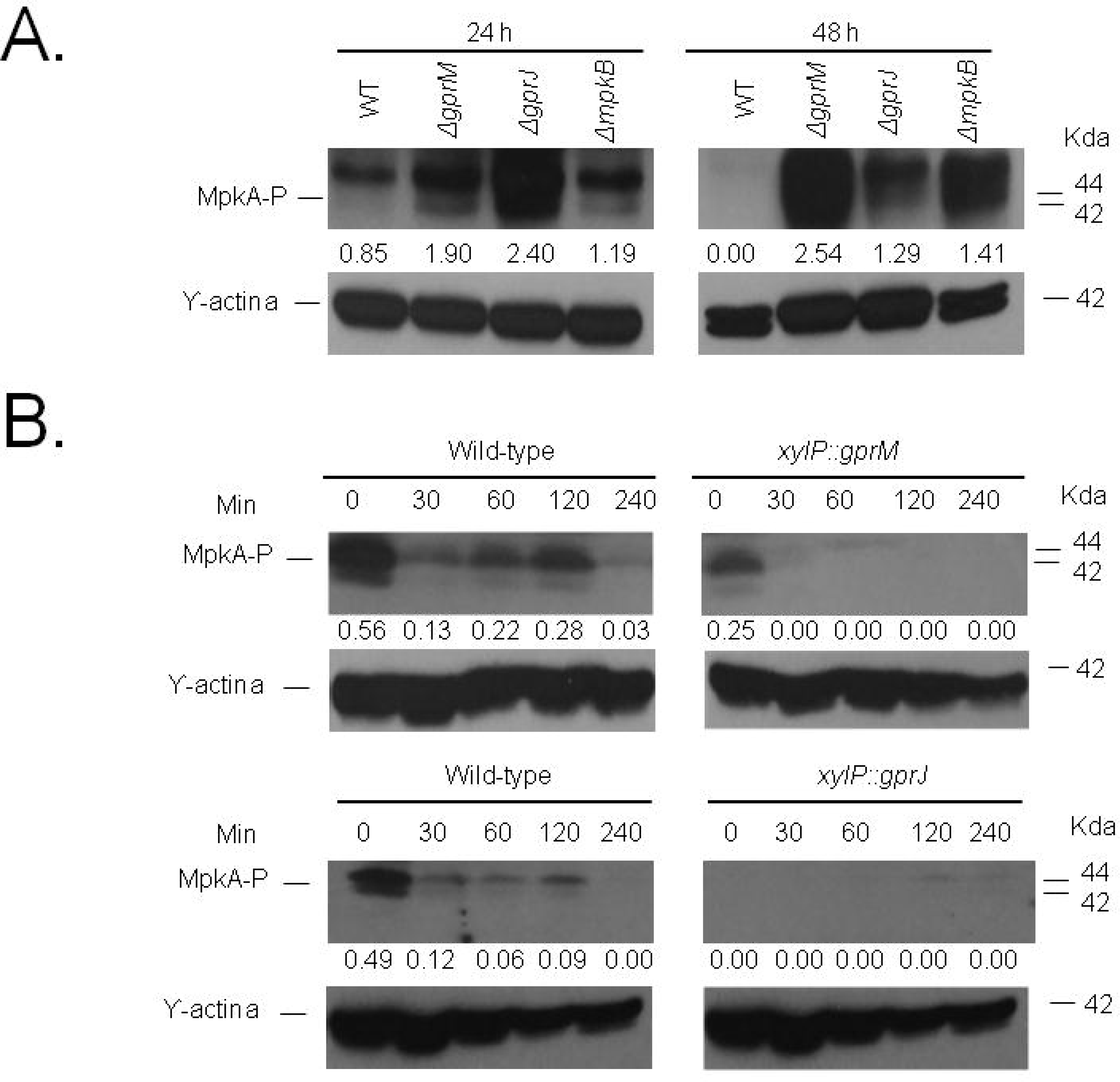
GprM and GprJ affect MpkA phosphorylation. A. Deletion strains were grown for 24 or 48 h in glucose minimal medium (GMM) and B. overexpression strains were grown in GMM before mycelia were transferred to xylose minimal medium for 30, 60, 120 and 240. Total cellular proteins were extracted, quantified and Western blots were run against phosphorylated MpkA (MpkA-P) using the anti-44/42 MpkA antibody. MpkA phosphorylation signals were normalised by cellular ϒ-actin using the anti-ϒ-actin antibody. Signal intensities were quantified using the ImageJ software, and ratios of MpkA-P to γ-actin were calculated.

To further investigate the role of GprM and GprJ in cell wall maintenance, we measured concentrations of cell wall polysaccharides and cell wall thickness in the respective deletion strains. Deletion of *gprM* and *gprJ* resulted in increased concentrations of cell wall glucosamine, glucose, galactose and N-acetylglucosamine; whereas the simultaneous deletion of *gprM* and *gprJ* restored cell wall sugar concentrations back to levels observed in the WT and complemented strains (Figure 5A-B). Transmission Electron Microscopy (TEM) analysis of Δ*gprM* and Δ*gprJ* hyphal germlings and conidia showed that their cell walls were significantly thicker than the cell walls of the WT and complemented strains (Figure 5C-D; Supplementary Table S2). Together these results suggest that GprM and GprJ regulate cell wall maintenance, likely through the MpkA CWI pathway. However, the simultaneous absence of both GPCRs activates an unknown compensatory signalling mechanism that maintains normal cell wall organization.

**Figure 5.**
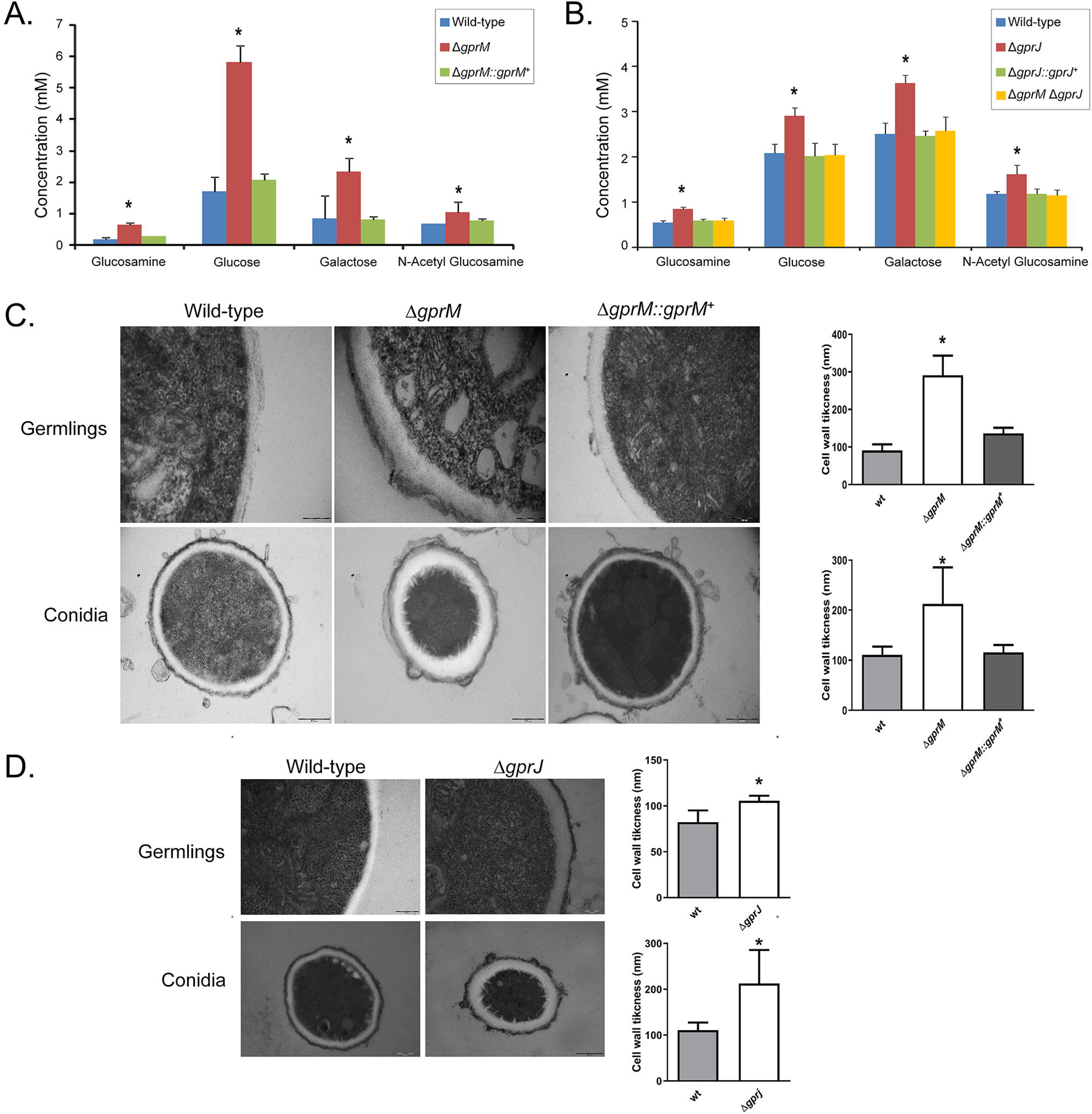
GprM and GprJ affect the cell wall organization. A.-B. Single deletion of *gprM* and *gprJ* results in increased concentrations of cell wall sugars. Strains were grown for 24 h in glucose minimal medium (GMM) before cell wall soluble fractions were prepared and high performance liquid chromatography (HPLC) was performed. Standard deviations present the average of six biological repetitions *p < 0.05 in a one-way ANOVA test in comparison to the wild-type strain. C.-D. Transmission electron microscopy of hyphal germlings and conida of the Δ*gprM* (C) and Δ*gprJ* (D) strains when grown for 24 h in GMM. Shown are representative pictures of hyphal and conidia cell wall sections as well as graphs showing the average cell wall thickness (nm) of 100 sections of different hyphal germlings or conidia (average of 4 sections per germling). Standard deviations present the average of 100 measurements with *p < 0.00001 in a one-tailed, paired t-test in comparison to the wild-type (WT) strain.

### GprM affects protein kinase A (PKA) activity

The cAMP-PKA pathway is activated in response to GPCR heterotrimeric G-protein signalling and regulates a variety of cellular processes, including fungal development and carbon source (e.g. glucose) utilization in *Aspergillus* species [7, 17]. Furthermore, the cAMP-PKA pathway is also important for sugar metabolism, carbohydrates that are build blocks for cell wall polysaccharide precursors in *A. fumigatus* [18]. We investigated whether cAMP-PKA signalling is activated by GprM and GprJ in the presence of glucose. PKA activity in the single and double mutants is comparable to the WT strain after 24 h growth in GMM, but Δ*gprM* and Δ*gprM* Δ*gprJ* have significantly reduced PKA activity after 48 h growth in GMM (Figure 6). Reduction of PKA activity in the Δ*gprM* Δ*gprJ* had no additive interaction, as PKA activity in the double mutant was mostly reminiscent of Δ*gprM*. Growth in the presence of glucose, and glucose transport were not significantly different in the single and double *gprM* and *gprJ* deletion strains than when compared to the WT strain (Supplementary Figure S2). Together, these results suggest that GprM, but not GprJ, activates the cAMP-PKA signaling pathway and that, in this case, the cAMP-PKA pathway is required for regulation of cellular processes that are not related to glucose metabolism.

**Figure 6.**
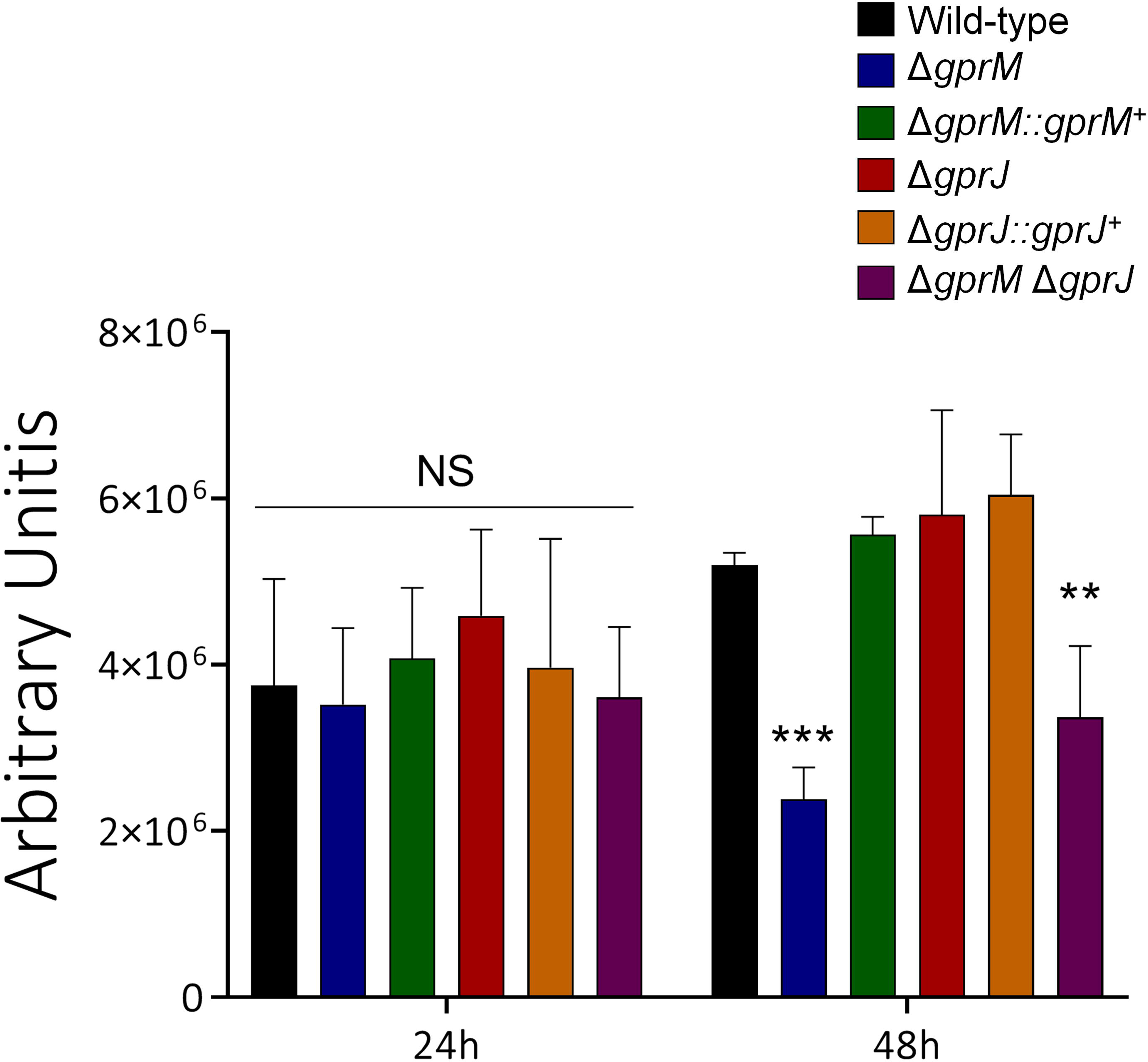
Protein kinase A (PKA) activity is reduced in the Δ*gprM* and Δ*gprM* Δ*gprJ* mutants. Strains were grown for 24 or 48 h in glucose minimal medium before total cellular proteins were extracted and PKA activity was measured. Standard deviations present the average of three biological replicates with **p <0.01 and ***p <0.001 (NS=non significant) in a two-way ANOVA followed by Bonferroni posttests.

### Transcriptional profiling of the *gprM* and *gprJ* loss of function and overexpression strains

To determine which genes are under the regulatory control of GprM and GprJ, we performed RNA-sequencing of the single deletion strains in GMM and of the overexpression strains in the presence of GMM and XMM. Differentially expressed genes (DEGs) were defined as those with a minimum log2 fold change of 1 [log2FC ≥ 1.0 and ≤ -1.0; *p*-value < 0.05; FDR (false discovery rate) of 0.05] when comparing the deletion or overexpression strains to the WT strain.

The WT, Δ*gprM*, and Δ*gprJ* transcriptional response was assessed after 24 h growth in GMM. In the Δ*gprM* and Δ*gprJ* strains, 70 and 238 genes were up-regulated respectively; whereas 64 and 21 genes were down-regulated respectively (Figure 7A). Of these, 66 and 11 genes were up- and down-regulated respectively in both deletion strains (Figure 7A). FunCat (https://elbe.hki-jena.de/fungifun/fungifun.php) enrichment analyses for the Δ*gprM* and Δ*gprJ* strains demonstrated a transcriptional up-regulation of genes coding for proteins involved in secondary metabolism, toxins, and metabolism of melanins (Figure 7B). FunCat analysis did not show any significant enrichment for the down-regulated genes probably due to the low number of DEGs. We then focused on the FunCat enrichment analysis of DEGs common to both strains and specific for each deletion strain. Most of the up-regulated DEGs shared between the mutants encode proteins involved in secondary metabolism (Figure 7C). The 172 DEGs specifically up-regulated in the Δ*gprJ* strains encoded proteins involved in secondary metabolism, metabolism of melanins, and non-ribosomal peptides (Figure 7C).

**Figure 7.**
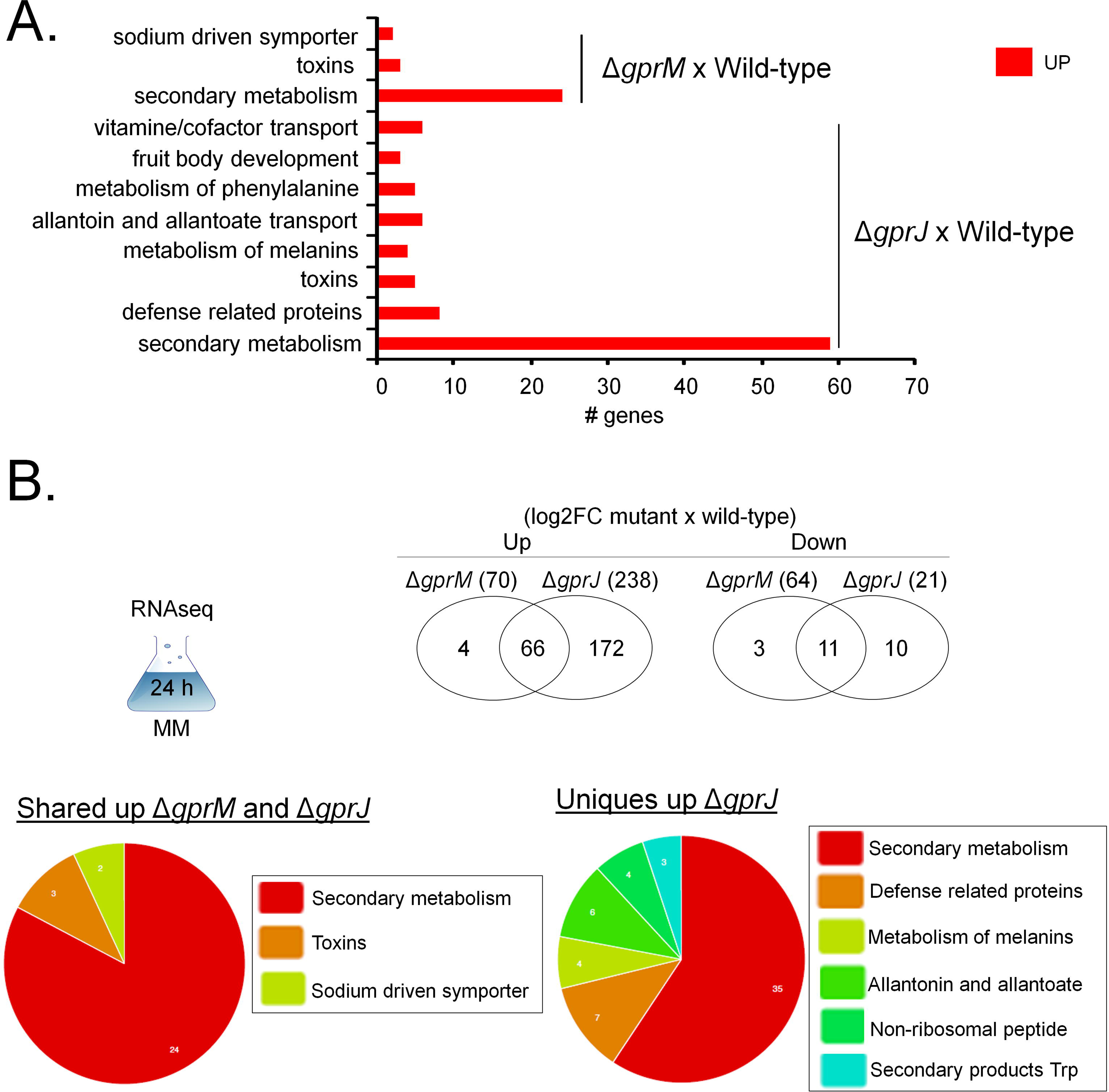
Functional characterisation (FunCat) of signficiantly differently expressed genes (DEGs) identified by RNA-sequencing in the Δ*gprM* and Δ*gprJ* strains. A. FunCat anlysis of DEGs up-regulated in the Δ*gprM* and Δ*gprJ* strains in comparison to the wild-type (WT) strain when grown for 24 h in glucose minimal medium (GMM). B. Venn diagrams showing the number of up- or down-regulated DEGs specific to each deletion strain and shared among the Δ*gprM* and Δ*gprJ* strains in comparison to the WT strain. In brackets is the total number of DEGs identified for each strain (log2FC = log2 fold-change). A diagram depicting the growth condition used for RNA-sequencing is also shown. In addition, FunCat analyses for up-regulated DEGs shared among the two deletion strains and for the Δ*gprJ* strain in comparison to the WT are shown as pie charts.

Next, we analysed the transcriptional profiles of GprM and GprJ in conditions of gene overexpression. The *xylP::gprM* and *xylP::gprJ* strains were grown for 16 h in GMM before being transferred to 1 % w/v xylose MM for 4 h (*xylP::gprM*) and 1 h (*xylp::gprJ*) to induce gene overexpression (Supplementary Tables S5 and S6). In the *xylp::gprM* and *xylp::gprJ* strains, 1436 and 40, and 1199 and 239 genes were significantly up-regulated and down-regulated, respectively (Figure 8A and Supplementary Tables S5 and S6). Of these, 7 and 58 genes were up- and down-regulated respectively in both overexpression strains (Figure 8A). FunCat enrichment analyses of the *xylp::gprM* strain showed a transcriptional up-regulation of genes encoding proteins involved in secondary metabolism and transport ATPases; and transcriptional down-regulation of genes coding for proteins involved in electron transport, secondary metabolism, transport facilities, C-compound and carbohydrate metabolism, and mitochondrion (Figure 8B). FunCat enrichment analysis of up-regulated DEGs in the *xylP::gprJ* could not be carried out due to the low number of DEGs. FunCat analysis of down-regulated DEGs in the *xylp::gprJ* strain showed enrichment of C-compound and carbohydrate metabolism and secondary metabolism enrichment (Figure 8C). We were unable to carry out FunCat analyses for the DEGs that are under the regulatory control of both GPCRs which was again due to the low number of DEGs.

**Figure 8.**
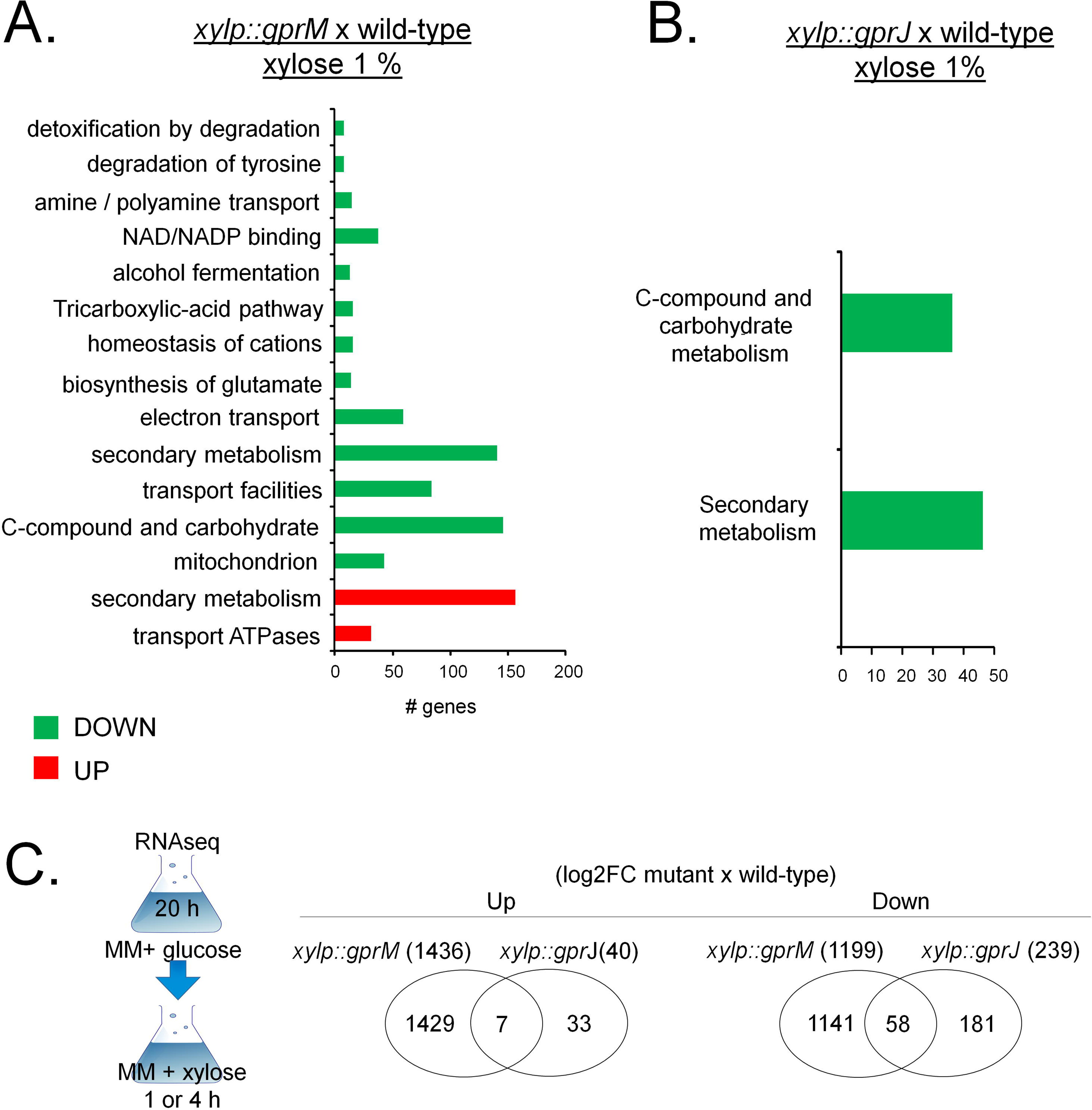
Functional characterisation (FunCat) of signficiantly differently expressed genes (DEGs) identified by RNA-sequencing in the *gprM* and *gprJ*-overexpressing strains. A. FunCat anlysis of DEGs up- and down-regulated in the *gprM* overexpressing strain in comparison to the wild-type (WT) strain when grown for 4 h in xylose minimal medium (XMM) after transfer from 24 h growth in glucose minimal medium (GMM). B. FunCat anlysis of DEGs down-regulated in the *gprJ* overexpressing strain in comparison to the WT strain when grown for 1 h in XMM after transfer from 24 h growth in GMM. C. Venn diagrams showing the number of up- or down-regulated DEGs specific to each deletion strain and shared among the Δ*gprM* and Δ*gprJ* strains in comparison to the WT strain (log2FC = log2 fold-change). In brackets is the total number of DEGs identified for each strain. A diagram depicting the growth condition used for RNA-sequencing is also shown.

Taken together, the data suggest that GprM and GprJ each regulate a unique set of genes and that not many genes are target to regulation by both GPCRs in the here tested conditions.

### GprM and GprJ are important for the regulation of the production of secondary metabolites (SMs)

We then focused on genes that were differentially expressed in our RNA-seq data and encoded SM-biosynthesis proteins, as we hypothesized that GprM and GprJ are involved in the regulation of SM biosynthesis. We also added the transcriptional profile of Δ*mpkB* when grown for 24 h in GMM to this analysis as a control (Figure 9; Supplementary Table S7). Visual inspection of DEGs encoding SM-related proteins, showed that biosynthetic gene clusters (BGC) of fumagillin, pyripyropene, fumigaclavine C, fumiquinazoline, melanin, and fumitremorgin were up-regulated in the Δ*gprM*, Δ*gprJ*, and Δ*mpkB* strains and down-regulated in the *xylp::gprM* and *xylp::gprJ* strains, suggesting that GprM and GprJ are important for the transcriptional regulation of SM BGCs (Figure 9).

**Figure 9.**
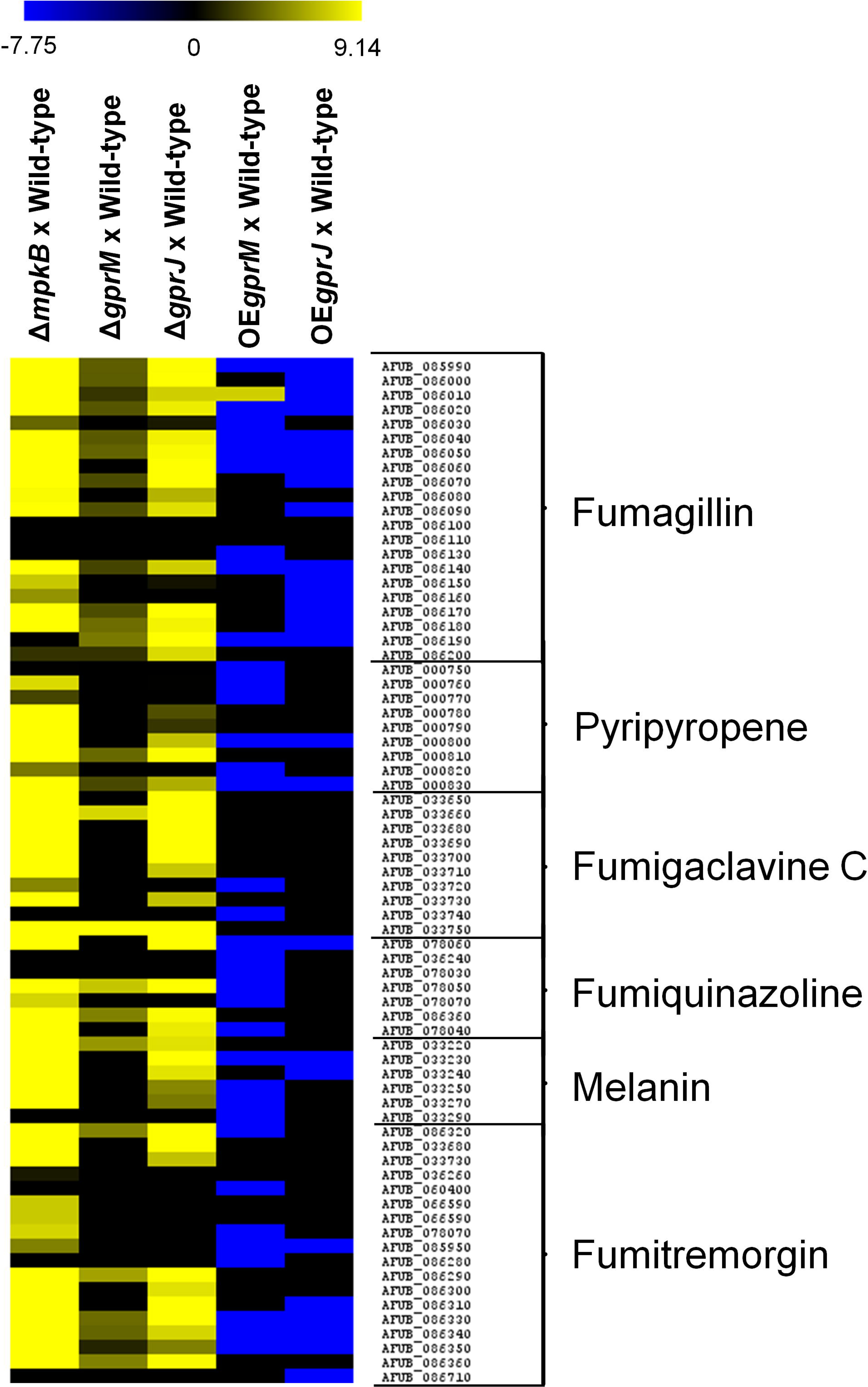
GprM and GprJ are important for the regulation of genes encoding proteins involved in secondary metabolite (SM) biosynthesis. Heat map depicting the log2 fold change (Log2FC) of differentially expressed genes (DEGs), as determined by RNA-sequencing, and encoding enzymes required for specific SM biosynthesis. Log2FC values are based on comparisons between the *mpkB* (control), *gprM*, *gprJ* deletion strains and the wild-type strain; as well as between the *gprM* and *gprJ* overexpressing (OE) strains and the wild-type strain. Heat map scale and gene identities are indicated. Hierarchical clustering was performed in MeV (http://mev.tm4.org/), using Pearson correlation with complete linkage clustering.

To confirm a role of both GPCRs in SM production, mass spectrometry (MS) analysis of supernatants of strains grown for 5 days in MM at 37 °C was carried out. MS analysis detected the presence of increased levels of fumagillin, pyripyropene, fumigaclavine C, fumitremorgin C and fumiquinazolines F and C in the Δ*gprM*, Δ*gprJ*, and Δ*mpkB* strains when compared to the WT strain (Figures 10A to 10G). These results are in agreement with the RNA-seq data. Since Δ*gprM*, Δ*gprJ*, and Δ*mpkB* strains have increased MpkA phosphorylation (Figure 4), we wondered whether the production of these SMs was also altered in the Δ*mpkA* strain (Figures 10A to 10G). The Δ*mpkA* strain had increased production of fumagillin and fumitremorgin in comparison to the WT strain, whereas concentrations of pyripyropene A, fumigaclavine C, and fumiquinazolines F and C were reduced in this strain (Figures 10A to 10G).

**Figure 10.**
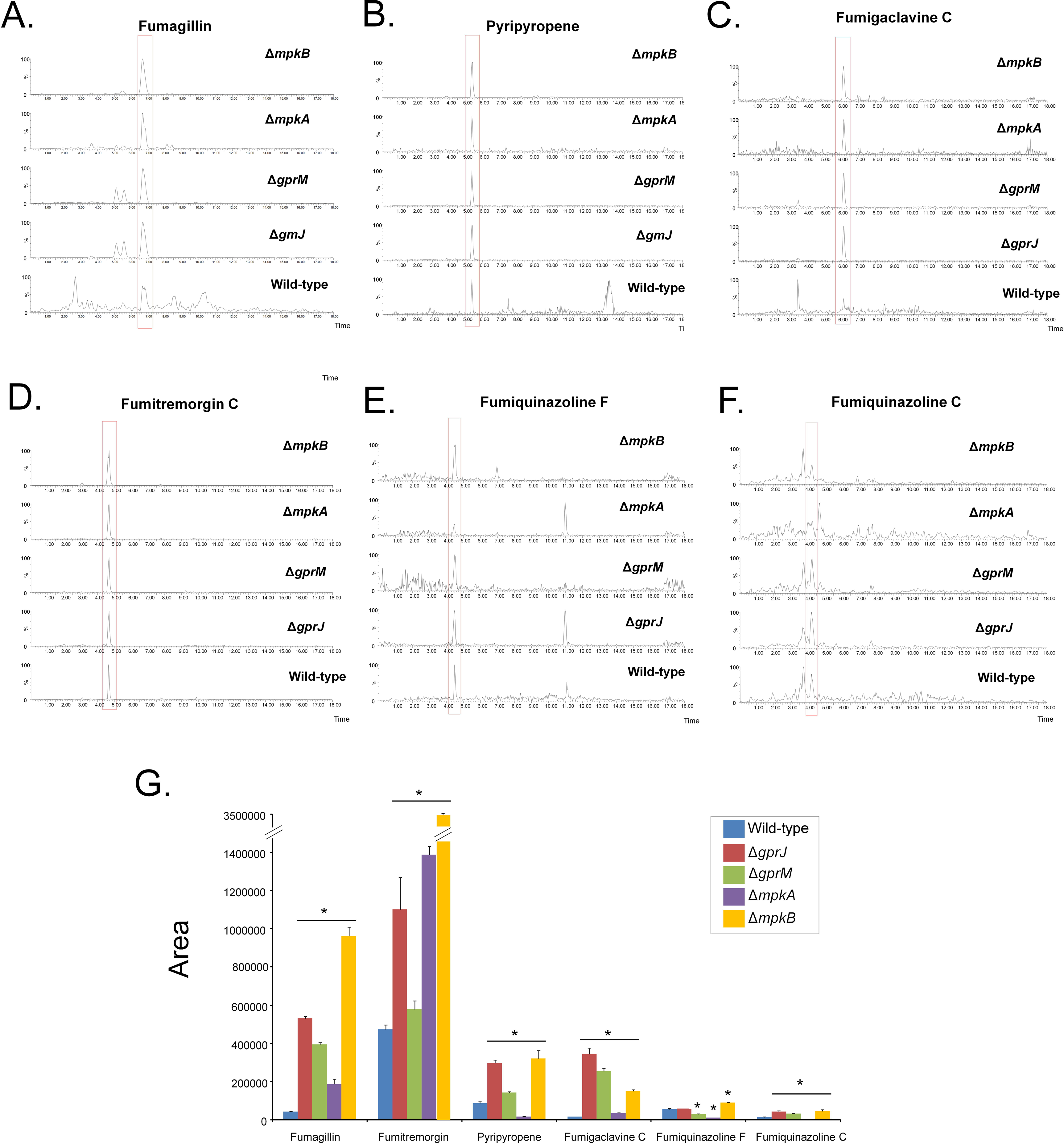
GprM and GprJ are involved in the repression of several secondary metabolites (SMs). SMs were extracted from culture supernatants of strains grown for 120 h in glucose minimal medium, before mass spectrometry (MS) was perfomed. A.-F. MS chromatograms of fumagillin (A), pyripyropene A (B), fumigaclavine C (C), fumitremorgin C (D), fumiquinazoline F (E), and fumiquinazoline C (F) generated for different strains. G. Quantitative comparison of the SMs identified in A-F between the deletion and the wild-type strains. Standard deviations represent the average of biological triplicates with *p < 0.05 in a two-tailed unpaired Student’s t-test.

These results indicate that GprM and GprJ negatively regulate the production of several secondary metabolites, and that this regulation occurs through the CWI pathway in a SM-dependent manner.

### Screening of an *A. fumigatus* transcription factor (TF) deletion library identifies a TF important for CWI

To identify effector components that act downstream of GprM and the CWI pathway, we screened our RNA-seq data for DEGs encoding TFs and found 44 genes of which 28 were up- and 16 were down-regulated in the *xylp::gprM* strain when grown for 4 h in XMM (Figure 11A). For this analysis, we focused on GprM because this receptor was shown to be important for PKA activity (Figure 6), a protein kinase that is involved in cell wall maintenance (de Assis *et al.*, 2018). Some of these genes were also modulated in the Δ*mpkB* and Δ*gprM* strains, but not in the Δ*gprJ* and *xylp::gprJ* strains (Figure 11A and Supplementary Tables S3 to S6, and S8). The TF deletion strains were grown on GMM supplemented with increasing concentrations of CR and CFW before radial diameter was measured and normalised by the drug-free, control condition (Figure 11B). The 1F3 (ΔAFUB_019790) strain was significantly more sensitive to CR and CFW (Figure 11B).

**Figure 11.**
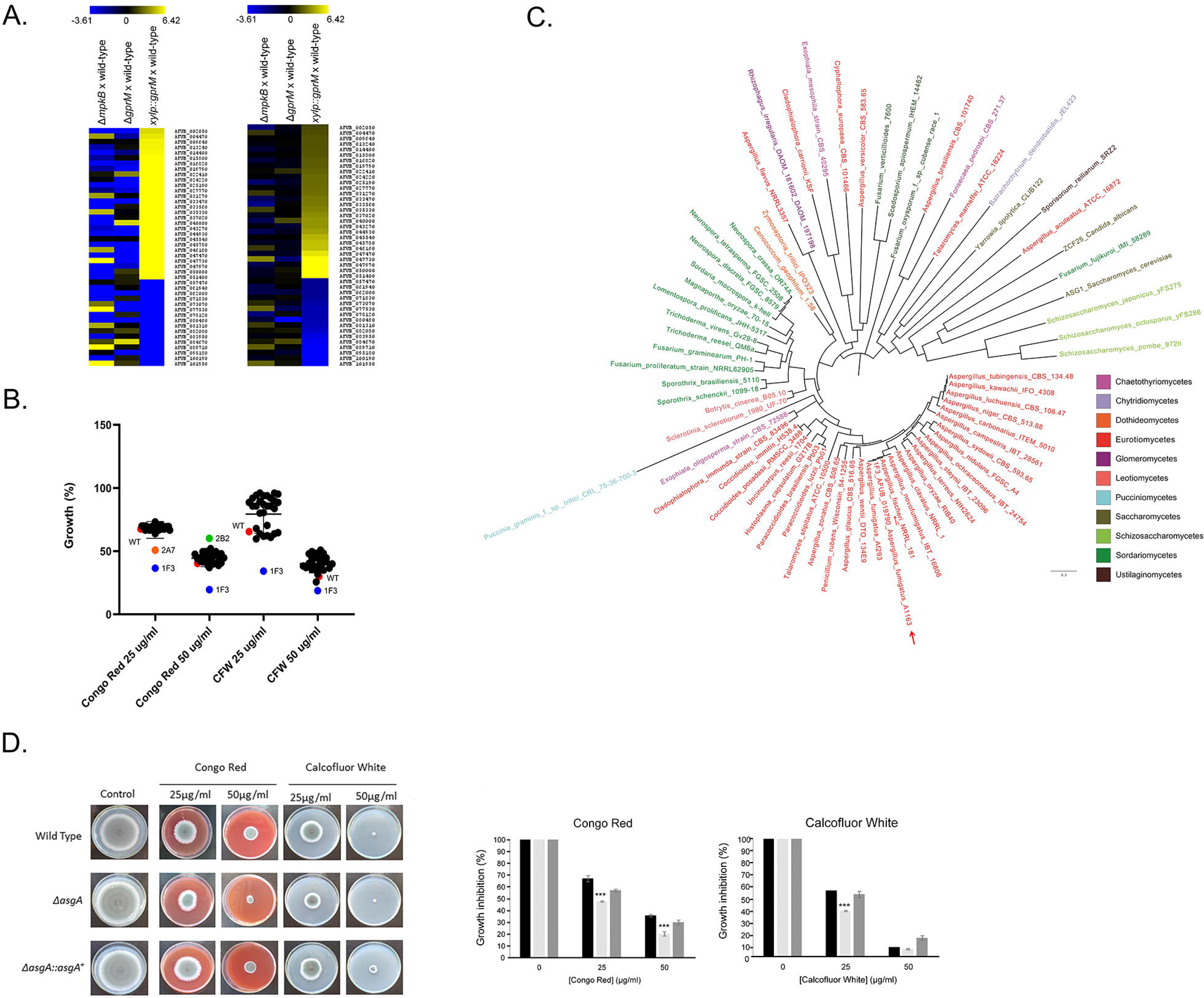
Identification of a TF involved in mediating the response to cell wall damage. A. Heat maps depicting the log2 fold change (Log2FC) of 44 differentially expressed genes (DEGs), as determined by RNA-sequencing, and encoding transcription factors (TFs) that were modulated in the *xylP::gprM* strain. Log2FC values are based on comparisons between the *mpkB* and *gprM* deletion strains and the *gprM* overexpressing (*xylP::gprM*) strain and the wild-type strain. Heat map scale and gene identities are indicated. Hierarchical clustering was performed in MeV (http://mev.tm4.org/), using Pearson correlation with complete linkage clustering. B. Identification of the TF AsgA as important for cell wall stress resistance. Graph showing the percentage of growth of 44 TF-deletion strains (named by simple numbers and letters) in the presence of different concentrations of the cell wall-damaging agents Congo red (CR) and calcofluor white (CFW). Growth % was calculated by dividing colony diameter in the presence of the drug by the colony diameter in the control, drug-free condition specific to each strain. The 1F3 strain, subsequently named Δ*asgA* strain, was significantly more sensitive to the cell wall-perturbing compounds. (C) The phylogenetic distribution of AsgA across fungal classes and genomes. Sequences were aligned through ClustalW implemented in the MEGA7 software (Kumar *et al.*, 2016). Phylogenetic analyzes were performed using the MEGA7 software, using the Neighbor-Joining method (Saitou and Nei, 1987) and 1000 bootstrap replications (Felsenstein, 1985) for each analysis. The phylogenetic tree was visualized using the Figtree program (http://tree.bio.ed.ac.uk/software/figtree/). The red arrow indicates the *asgA* gene. (D) Confirmation of AsgA-mediated sensitivity to CR and CFW. Strains were grown from 10^5^ spores for 5 days at 37 °C on glucose minimal medium supplemented with increasing concentrations of CR and CFW before pictures (left panel) were taken and radial diameter (right panel) of biological triplicates was measured. Standard deviations represent the average of thre biological replicates with ***p < 0.0005 in a one-tailed, paired t-test in comparison to the wild-type (WT) strain.

AFUB_019790 encodes a putative transcription factor of 895 amino acids with a molecular weight of 99.8 kDa; a GAL4-like Zn(II)2Cys6 (or C6 zinc) binuclear cluster DNA-binding domain (SM00066, https://smart.embl.de/smart; between amino acids 63 to 107) and a fungal specific transcription factor domain (SM000906, https://smart.embl.de/smart; between amino acids 416 to 489). AFUB_019790 has high sequence similarity at the protein level with *S. cerevisiae* Asg1p (24 % identity, 40 % similarity, e-value: 8e-33), a transcriptional regulator predicted to be involved in stress responses, as *ASG1* deletion strains have a respiratory deficiency, increased CFW sensitivity and slightly increased cycloheximide resistance [19]. Accordingly, we have named AFUB_019790 as *asgA*. Phylogenetic analysis of AsgA across fungal species representing the 13 different taxonomic classes or subphyla within Dikarya (www.fungidb.org), revealed that orthologues were largely distributed in the Pezizomycotina [Eurotiomycetes (Chaetothyriomycetes), Sordariomycetes, Leotiomycetes, and Dothideomycetes], Saccharomycotina (Saccharomycetes), Taphrinomycotina (Schizosaccharomycetes), Ustilaginomycotina (Ustilaginomycetes), P u c c i n i o m y c o t i n a (Pucciniomycetes), Glomeromycotina (Glomeromycetes), and Chytridiomycota (Chytridiomycetes; Figure 11C and Supplementary Table S8). AsgA orthologues are also present in other important fungal pathogens, such as *Botrytis cinerea, Coccidioides spp*, *Sporothrix spp*, *Paracoccidioides spp*, *Fusarium spp*, and *Ajellomyces capsulatus* (Supplementary Table S8).

We constructed the *asgA* complemented strain and confirmed that deletion of this TF-encoding gene resulted in increased sensitivity to CR and CFW when compared to the wild-type and complemented strains (Figure 11D).

### The Δ*gprM* and Δ*gprJ* mutants have reduced virulence in the *Galleria mellonella* insect model of invasive aspergillosis

We wondered whether GprM and GprJ are important for *A. fumigatus* virulence, since they are involved in cell wall maintenance, which is crucial for pathogenicity [20]. *G. mellonella* larvae (n = 10 for each strain) were infected with the WT, *gprM* and *gprJ* deletion and complementation strains and survival was assessed over a time period of 10 days (Figure 12A to 12C). The wild-type, Δ*gprM::gprM^+^*, and Δ*gprJ::gprJ^+^* strains caused 100 and 90 % mortality after 10 days post-infection (p.i.), respectively (Figures 12A and 12B). In contrast, the Δ*gprM* and Δ*gprJ* strains caused 40 and 50 % mortality rate 10 days p.i., which was statistically (p<0.001, Mantel-Cox and Gehan-Brestow-Wilcoxin tests) different from the WT and complemented strains (Figures 12A and 12B). The Δ*gprM* Δ*gprJ* strain caused 70 % mortality at 10 days p.i., which was not statistically different from the WT and complemented strains (p<0.001; Figure 12C).

**Figure 12.**
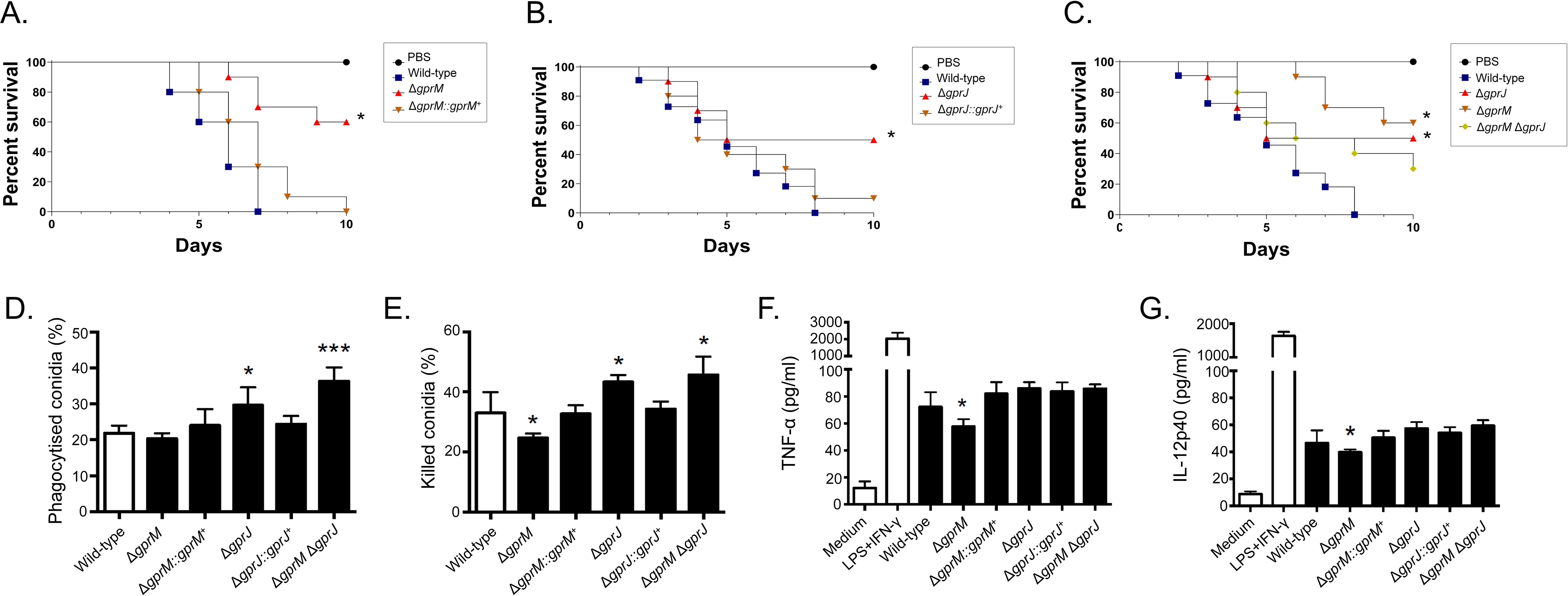
GprM and GprJ are important for *A. fumigatus* virulence in *G. mellonella* wax moth. A.-C. Survival curves of *G. mellonella* larvae (n = 10/fungal strain) infected via injection with 10^6^ conidia of the *gprM* and *gprJ* single and double deletion strains. Larval survival was monitored over a time period of 10 days. D.-E. Bone marrow-derived murine C57BL/6 macrophages (BMDMs) phagocytize (D) and kill (E) a higher number of Δ*gprJ* and Δ*gprM* Δ*gprJ*, conidia *in vitro*, whereas they kill less Δ*gprM* conidia. F.-G. Concentrations of the tumor necrosis factor alpha (TNF-α) and interleukin (IL)-12p40 cytokines in BMDMs infected with live-resting (LR) conidia, show that Δ*gprM* conidia elicit a reduced inflammatory response. Positive control: LPS+IFN-λ= Lipopolysaccharide+interferon λ. Standard deviations represent the average of three biological replicates and all the strains were compared with the wild-type and complemented strains (*p-value < 0.05 in a two-tailed, unpaired student t-test).

Since, deletion of *gprM* and *gprJ* caused alterations in cell wall composition (Figure 5), we hypothesized that this could influence the host immune response. Macrophages contribute to innate immunity, fungal clearance and the generation of a pro-inflammatory response during *A. fumigatus* infection [21]. The capacity of bone marrow-derived macrophages (BMDMs) to phagocytize and kill conidia derived from the WT, Δ*gprM*, Δ*gprM::gprM*^+^, Δ*gprJ*, Δ*gprJ::gprJ*^+^, and Δ*gprM* Δ*gprJ* strains was assessed.

Conidia from the Δ*gprJ* and Δ*gprM* Δ*gprJ* strains are significantly more phagocytosed than conidia from the WT and Δ*gprJ::gprJ* strains, whereas no differences in phagocytosis were observed for the other strains (Figure 12D). In agreement with the increased phagocytosis, conidia from the Δ*gprJ* and Δ*gprM* Δ*gprJ* strains were killed significantly more than when compared to the WT and gprJ complemented strains (Figure 12E). In addition, the Δ*gprM* conidia are killed less than conidia from the WT and Δ*gprM::gprM*^+^ strains (Figures 12E). Interestingly, there is a reduced production of the cytokines TNF-α and IL12-p40 in the Δ*gprM* strain when compared to the WT and Δ*gprM::gprM*^+^ strains (Figures 12F and 12G). There are no differences of these cytokines in the Δ*gprJ* and Δ*gprM* Δ*gprJ* strains when compared to the WT and Δ*gprJ::gprJ^+^* strains (Figures 12F and 12G).

Taken together, these results indicate that Δ*gprM* and Δ*gprJ* are important for *A. fumigatus* virulence in *G. mellonella* whereas the double mutant is not; suggesting the existence of an unknown compensatory signalling mechanism that promotes virulence.

## Discussion

Fungi can sense their environment through the presence of membrane-associated GPCRs, which activate downstream signalling cascades and allow the fungus to appropriately respond to environmental cues. In the case of human pathogenic fungi, such as *A. fumigatus*, these membrane receptors are predicted to be crucial for mammalian host infection, and are suggested targets for the development of anti-fungal drugs [6, 7, 22]. In this work, we have characterized two *A. fumigatus* GPCRs and have shown their importance for several virulence determinants such as the CWI pathway, secondary metabolite production, and infection.

To date, the genome of *A. fumigatus* is predicted to encode 15 GPCRs which remain largely uncharacterized with the exception of a couple of studies which functionally characterized the three GPCRs GprC, GprD and GprK [10, 11]. Furthermore, the great number of GPCRs in fungal genomes makes their study difficult because the majority of single deletion mutants of GPCR genes do not show a clear phenotype due to the existence of redundancy between them [23-25]. In agreement, deletion of *gprM* and *gprJ* resulted in two strains with increased melanin production in supernatants of liquid cultures. A function for GprM in repressing DHN-melanin production has previously been described [12]. Furthermore, GprM physically interacts with the Gα protein GpaA, with deletion of *gpaA* resulting in a strain with increased DHN-melanin production [12]. It was suggested that GprM-GpaA signalling occurs through the MAPK MpkB, resulting in the regulation of SM production, including the biosynthesis of DHN-melanin [12]. Recently, this signaling cascade consisting of the three kinases SteC, MkkB, and MpkB, as well as the SteD adaptor protein and the HamE scaffold was shown to be critical for the regulation of fungal development and secondary metabolism [26]. In this study, we identified an additional GPCR, GprJ, that may also be involved in controlling DHN-melanin production. We are currently performing studies to determine whether GprJ also physically interacts with GpaA and whether it activates the MpkB signalling pathway. It is likely that GprJ at least participates in the regulation of the MpkB pathway, as GprJ was shown to regulate phosphorylation of MpkA, the MAPK of the CWI pathway which indirectly interacts with MpkB [12].

Furthermore, this study showed a genetic interaction between *gprM* and *gprJ* which appears to be highly complex. In the Δ*gprM* and Δ*gprJ* strains, similar phenotypes are observed for cell wall polysaccharide concentrations and virulence in *G. mellonella*. Simultaneous deletion of both GPCR-encoding genes resulted in a strain with a cell wall and virulence similar to the WT strain, but not in the MpkA phosphorylation profile, suggesting that the absence of both receptors activates unknown compensatory signalling mechanisms to promote normal MpkA-independent cell wall organization and virulence. Transcriptional profiling of both single deletion strains showed that few genes are targeted by both GPCRs, suggesting that GprM- and GprJ-mediated signalling differs significantly and may occur via different pathways. This is supported by the observation that PKA activity is decreased in the Δ*gprM* and Δ*gprM* Δ*gprJ* strains but not in the Δ*gprJ* strain. Restauration of WT phenotypes upon deletion of two genes encoding GPCRs has already been observed in other fungi. In *A. nidulans*, the addition of glucose after carbon starvation resulted in a burst in cAMP, a response that was absent in the Δ*gprH*, Δ*gprI* and Δ*gprM* single deletion strains [17]. In contrast, the simultaneous absence of multiple GPCRs, in the double or triple deletion strains, resulted in the recovery of the cAMP burst after addition of glucose [17]. These studies show that phenotypes observed in single deletion strains can cancel each other out and that this may also be the case for the Δ*gprM* Δ*gprJ* strain for certain conditions as these GPCRs may regulate signalling pathways in an opposite manner.

It is clear though from our study that GprM and GprJ, as well as downstream signalling events, are involved in SM production in *A. fumigatus*. This is in agreement with previous studies which showed that the deletion of the G-protein α subunit, GpaB and the adenylate cyclase AcyA decreased DHN-melanin production in *A. fumigatus* [27] whilst overexpression of the protein kinase A catalytic subunit PkaC1 induced the expression of the DHN-melanin cluster genes [28]. Genes encoding proteins required for the biosynthesis of SMs were shown to be under the transcriptional control of GprM and GprJ. Indeed, GprM and GprJ are involved in the repression of SM secretion, as the concentrations of secreted SMs such as fumagillin, pyripyropene, fumigaclavine C, fumiquinazoline, melanin, and fumitremorgin were significantly reduced in the Δ*gprM* and Δ*gprJ* strains. This is in agreement with a study that showed that GprM negatively regulates DHN-melanin production [12]. GprM- and GprJ-mediated regulation of SM biosynthesis may occur via different MAPK signalling pathways, including MpkB (as discussed above) and MpkA. Extracellular concentrations of pyripyropene, fumigaclavine C, fumiquinazoline, and melanin were not detected in the *mpkA* null mutant while fumagillin and fumitremorgin are negatively regulated by MpkA, suggesting that regulation through MpkA occurs in a metabolite-dependent manner. MpkA was indeed shown to be important for the regulation of genes encoding proteins required for gliotoxin, pyomelanin, pseurotin A, and siderophore biosynthesis [29]. The Δ*mpkA* strain showed reduced DHN-melanin and gliotoxin production but had increased production of siderophores during iron starvation conditions [16, 29-31]. Regulation of SM production is therefore likely to occur via various signalling pathways which are activated upon sensing many different environmental cues. Adding to this complexity is the existence of cross-talk between different GPCR downstream signalling pathways which has been shown to regulate a number of virulence determinants, including SM production and CWI, in *A. fumigatus* [12, 18]. In the plant pathogen *A. flavus*, deletion of the 15 GPCR-encoding genes resulted in strains with one or more defects in growth, development, aflatoxin production, response to carbon sources, nitrogen sources, stress agents, and lipids; suggesting cross-talk between downstream signalling pathways [23]. It is tempting to suggest that each extracellular signal results in the activation of a common and “core” signalling pathway which interacts with a specific set of proteins to fine tune the cellular response to the environmental cue.

Furthermore, GprM and GprJ were shown to regulate *A. fumigatus* CWI, a process that is likely to occur through the MAPK MpkA. Phosphorylation of MpkA is synonymous with CWI pathway activation [16]. Deletion of *gprM* and *gprJ* resulted in an accumulation of phosphorylated MpkA and thus a “hyperactive” CWI pathway. This activation of CWI signalling may be the cause for the observed accumulation of cell wall polysaccharides and significantly increased cell wall thickness. In contrast, overexpression of *gprM* and *gprJ* reduced MpkA phosphorylation and inactivated the CWI pathway. As a consequence, cell wall thickness was significantly reduced and strains were sensitive to cell wall-interfering agents. Although the CWI pathway is well characterised in *A. fumigatus*, less is known about MpkA downstream targets. A notable exception is the TF RlmA, which acts downstream of MpkA, is involved in MpkA phosphorylation and required for a normal cell wall organization and for vegetative growth [32]. Recently, the TF ZipD together with the calcium-dependent TF CrzA, were shown to also influence MpkA phosphorylation levels and to be important for cell wall maintenance through regulating the expression of glucan and chitin biosynthetic genes in a calcium-dependent manner [33, 34]. To identify additional TFs that act downstream of the GprM-MpkA signalling pathway, we selected 44 TF deletion strains, whose expression was significantly altered in our *gprM* overexpression RNA-sequencing data, and assessed their growth in the presence of cell wall interfering agents. The Δ*asgA* was significantly more sensitive to cell wall perturbing agents. In agreement with its role in cell wall maintentance, AgsA was shown to have increased phosphorylation upon CR exposure [35]. We are currently performing additional work to decipher the relationship between AgsA and MpkA in the presence of cell wall stress.

In summary, this study functionally describes two *A. fumigatus* GPCRs and significantly contributes to our understanding of these types of signaling modules in this opportunistic fungal pathogen. Furthermore, this study also shows the complexity and cross-talk that underlies fungal GPCR-signaling in regulating important virulence traits. As fungal GPCRs are poorly conserved in animals and plants, and regulate virulence determinants in pathogenic fungi, they continue to be interesting targets for the development of novel anti-fungal drug and combinational therapies.

## Materials and Methods

### GPCRs among *Aspergillus* species

To identify GPCRs across *Aspergillus* species, we used a best-blast-hit based approach [36]. More specifically, we first obtained *Aspergillus* GPCR sequences based on a previously published list [9]. Using the *Aspergillus* GPCR protein sequences, we used Blast+, v2.3.0 [36], to identify GPCRs among 52 publicly available and annotated *Aspergillus* genomes from NCBI’s Genbank database (download date 2018-06-22). GPCR sequences were aligned and trimmed as described elsewhere [37]. Briefly, sequences were aligned using Mafft, v.7.402 [38], (parameters: --bl 62, --op 1.0, --maxiterate 1000, --retree 1, --genafpair). Nucleotide sequences were then threaded onto the protein alignment using Biopython, v1.7 [39], and trimmed using trimAl with the ‘gappyout’ parameter [40]. The resulting trimmed alignment was used to infer the evolutionary relationships among GPCRs using IQ-TREE, v1.6.1 [41]. Bipartition support was assessed using 5,000 ultrafast bootstrap approximations [42]. Bipartitions with less than 85% ultrafast bootstrap approximation support were collapsed.

### Strains and media

Strains were grown at 37°C in either complete medium (YAG) [2% (w/v) glucose, 0.5% (w/v) yeast extract, trace elements] or minimal medium [1% (w/v) glucose, nitrate salts, trace elements, pH 6.5]. Solid YAG and MM were the same as described above except that 1.7% (w/v) or 2% (w/v) agar was added. Trace elements, vitamins, and nitrate salts compositions were as described previously [43].

For phenotypic characterization, solid medium plates were inoculated with 10^4^ conidia per strain and left to grow for 120 h at 37°C. Radial growth was expressed by the ratio of colony radial diameter in the stress condition by colony radial diameter in the control (no stress) condition. All the strains used in this work are described in the Supplementary Table S1.

### Enzymatic assays

PKA activity (PepTag® Non-Radioactive Protein Kinase Assay kit V5340, Promega) was carried out according to the manufacturer’s instructions. Extracellular glucose concentrations were quantified as described previously [17].

### Aspergillus fumigatus strains

Fungi strains and plasmids used in this study are listed in Supplementary Table S1. In addition, all primers used in this work are listed in the Supplementary Table S9.

The *gprJ* (Afu1g06840) gene was deleted by gene replacement with the *ptrA* gene from *A. oryzae* using homologous recombination [44] generating the Δ*gprJ* strain. Briefly, the 5’ and 3’ flanking regions of the *gprJ* gene were amplified from *A. fumigatus* genomic DNA (gDNA) using specific primers (P1-P2) and (P3-P4), respectively (Supplementary Table S9). These were subsequently fused by using *S. cerevisiae in vivo* recombination [45] to the *ptrA* gene, which confers resistance to pyrithiamine, and which was PCR amplified from plasmid pSK275 (primers P5/P6). The construct was used to transform the *A. fumigatus* Δ*akuB*^KU80^ wild-type strain via homologous recombination [46]. Mutants were selected and purified on MM containing 1 µg ml^-1^ pyrithiamine and confirmed with PCR using a primer (P29) annealing externally to the integrating cassette (Supplementary Figure S3).

The Δ*gprJ* strain was cultivated on complete medium supplemented with uridine, uracil and 0.75 mg ml^-1^ of 5-fluoroorotic acid to obtain a Δ*gprJ pyrG^−^* strain. The *ΔgprJ* pyrG^-^ strain was confirmed by PCR and used to generate the *gprJ* complementing strain, the Δ*gprJ* Δ*gprM* and Δ*gprJ* Δ*pksP* double mutants. Deletion cassettes containing the *pyrG* selectable marker were generated via yeast-mediated recombination as previously described [47]. Briefly, 1-kb fragments from the gene 5’ and 3’ flanking regions were amplified with the corresponding primer pairs [primers P13 through P16 for *gprM* gene (Afu7g05300) and P31 through P34 for *pksP* (Afu2g17600)]. The *pyrG* gene fragment was PCR amplified from the plasmid pCDA21 with primers P11 and P12. All fragments were co-transformed into the *Saccharomyces cerevisiae* SC9721 strain along with the linearized plasmid pRS426. Deletion cassettes were finally amplified from yeast genomic DNA with corresponding 5’ Fw and 3’ Rv primers and used to transform the auxotroph Δ*gprJ pyrG^−^* strain. Mutants were selected and purified in MM and were checked by PCR using external primers (P29 for *gprM* and P35 for *pksP*). To complement the Δ*gprJ* strain, a complementation cassette containing the *gprJ* ORF as well as the *pyrG* selectable marker was constructed as described above and used to transform the Δ*gprJ pyrG^−^* strain. Primers P8 and P9, containing homologous tags to the *pyrG* gene, were used to generate the cassette. The complementing cassete was transformed into the *ΔgprJ* pyrG^-^ strain and positive *A. fumigatus* complementing candidates were selected, purified through three rounds of growth on plates, submitted to gDNA extraction and confirmed by PCR (Supplementary Figure S3).

The overexpressing *xylp*::*gprJ* strain containing the *gprJ* gene N-terminally tagged to the xylose reductase promoter (*xylp*) and C-terminally tagged to HA plus the TRPC terminator and the *pyrG* auxotrophic marker was generated by homologous integration. The final cassette contained the following fragments: 5’-UTR (P7/P23), *xylp* (P17/P18), *gprJ* ORF without the stop codon (P21/P22), 3xHA-*trpC*-*pyrG* (P19/P20), and 3’-UTR (P24/P25). Primer pairs used for PCR amplification of each DNA fragment are indicated in parentheses. Fragments *xylp* and 3xHA-*trpC*-*pyrG* were PCR amplified from the plasmids pYES-pXyl-hph-devR and pOB430, respectively. All fragments were co-transformed into *S. cerevisiae* SC9721 and the resulting cassette was used to transform the Δ*akuB*^KU80^ *pyrG^−^* strain. Genomic integration of the cassette was verified by PCR (Supplementary Figure S3) and the functionality of the construction was assayed by real-time PCR (Primers P26/P27) (Supplementary Figure S3).

Complementation of the *asgA* gene (Afu2g02690) was achieved via co-transformation. A fragment containing the *asgA* gene flanked by 1-kb 5’ and 3’-UTR regions was PCR amplified (Primers P35/P36). This DNA fragment was transformed into the Δ*asgA* strain along with the plasmid pPTR1 (TaKaRa Bio) containing the pyrithiamine resistance *ptrA* gene. Mutants were selected on MM supplemented with 1µg ml^-1^ pyrithiamine and confirmed with PCR using the external primer (P37).

### Generation, validation, and screening of transcription factor knockout mutants

The transcription factor knockout mutant collection was generated in the *A. fumigatus* strain A1160 (a derivative of CEA17, ΔKu80 pyrG^+^) according to [48]. The most sensitive and resistant mutants to CR (congo red) and CFW (calcofluor white) were identified by growing them on MM+CR and MM+CFW for 5 days at 37 °C; they were retested and purified for further characterization.

### RNA sequencing, cDNA synthesis and real-time PCR

For RNA sequencing, *A. fumigatus* conidia (5×10^7^) from the CEA17, Δ*gprM*, and Δ*gprJ* strains were inoculated in triplicate in liquid MM and cultured for 24 h at 37°C. For the *xylp::gprM* or *xylp::gprJ* strains, the conidia were inoculated in MM+1% glucose, grown for 24 h at 37 °C and then transferred to MM+1 % xylose for 4 or 1 h respectively. Mycelia were harvested, frozen and ground in liquid nitrogen. Total RNA was extracted using Trizol (Invitrogen), treated with RQ1 RNase-free DNase I (Promega) and purified using the RNAeasy Kit (Qiagen) according to manufacturer’s instructions. RNA from each treatment was quantified using a Qubit fluorometer and analyzed using an Agilent 2100 Bioanalyzer system to assess the integrity of the RNA. All RNA had an RNA Integrity Number (RIN) between 7.0 and 9.5.

Illumina TruSeq Stranded mRNA Sample Preparation kit was used to construct cDNA libraries following manufacturer’s instructions. Libraries were sequenced (2×100bp) on the Brazilian Bioethanol Science and Technology Laboratory’s (CTBE) NGS sequencing facility using a HiSeq 2500 instrument, generating approx. 11×10^6^ fragments per sample. Short reads were submitted to the NCBI’s Short Read Archive under the Bioproject ID: PRJNA634102.

Obtained fastq files were quality checked with FastQC (http://www.bioinformatics.babraham.ac.uk/projects/fastqc/) and cleaned (quality trim, adaptor removal and minimum length filtering) with Trimmomatic [49]. Ribosomal RNA was removed using SortMeRNA [50]. High quality RNA-seq reads were mapped against the *A. fumigatus* genome using the CLC Genomics Workbench software (CLC bio - v4.0; Finlandsgade, Dk) using the following parameters: mapping settings (minimum length fraction = 0.7; minimum similarity fraction = 0.8, and maximum number of hits for read = 1) and alignment settings (minimum distance = 180 and maximum distance = 1000). All samples achieved saturation of known exon-exon junctions. Reproducibility among biological replicates was assessed by exploring a principal component analysis (PCA) plot of the top 500 genes have the largest biological variation between the libraries, and by pair-wise measuring the Pearson correlation among the replicates over the whole set of genes. In order to assess transcript abundance exonic reads were counted in a strand-specific way using the featureCounts function from Rsubread Bioconductor package [51]. Calling of differentially expressed genes was carried-out using DESeq2 (70) using as threshold adjusted p-value < 0.01 [52].

For quantitative real-time PCR analysis, 1 x 10^7^ spores of the wild-type, *xylp::gprM* and *xylp::gprJ* strains were inoculated in 50mL of liquid MM supplemented with 1% glucose and grown at 37°C. After 24h of growth mycelia were transferred to liquid MM supplemented with 1 % xylose for up to 4h. Then, mycelia were frozen in liquid nitrogen and the total cellular RNA was extracted using TRIZOL (Invitrogen) according to manufacturer’s instructions. Total RNA was purified (RNeasy^®^ Mini Kit - Qiagen), according to manufacturer’s instructions and the quality of the RNA was assayed using the Agilent Bioanalyser 2100 (Agilent technologies). RNA was reverse transcribed to cDNA using the ImProm-II™ Reverse Transcription System (Promega) and the synthesized cDNA was used for real-time analysis using the SYBR Green PCR Master Mix kit (Applied Biosystems) in the the ABI 7500 Fast Real-Time PCR System (Applied Biosystems, Foster City, CA, USA). Primer sequences are listed in Supplementary Table S9.

### Immunoblot analysis

To assess the phosphorylation status of MpkA, fresh harvested conidia (1×10^7^) of the wild-type and deletion strains were inoculated in 50 ml liquid MM at 37°C for 24 h (180 rpm). For the overexpression strains, the wild-type and the mutant strains were grown at 37 °C for 24 h in MM and transferred for MM+1 % xylose at 37 °C for 30 to 240 min. Mycelia were ground in liquid nitrogen with pestle and mortar. For protein extraction, 0.5 ml lysis buffer containing 10% (v/v) glycerol, 50 mM Tris–HCl pH 7.5, 1% (v/v) Triton X-100, 150 mM NaCl, 0.1% (w/v) SDS, 5 mM EDTA, 50 mM sodium fluoride, 5 mM sodium pyrophosphate, 50 mM β-glycerophosphate, 5 mM sodium orthovanadate, 1 mM PMSF, and 1X Complete Mini® protease inhibitor (Roche Applied Science) was added to the ground mycelium. Extracts were centrifuged at 20,000 g for 40 minutes at 4°C. The supernatants were collected and the protein concentrations were determined using the Bradford assay (BioRad). Fifty µg of protein from each sample were resolved in a 12% (w/v) SDS–PAGE and transferred to polyvinylidene difluoride (PVDF) membranes (Merck Millipore). The phosphorylated fractions of the MAP kinase, MpkA, were examined using anti-phospho p44/42 MAPK antibody (Cell Signaling Technologies) following the manufacturer’s instructions using a 1:1000 dilution in TBST buffer (137 mM NaCl, 20 mM Tris, 0.1% Tween-20). Anti-γ-actin was used to normalize the protein loading. Primary antibody was detected using an HRP-conjugated secondary antibody raised in rabbit (Sigma). Chemoluminescent detection was achieved using an ECL Prime Western Blot detection kit (GE HealthCare). To detect these signals on blotted membranes, the ECL Prime Western Blotting Detection System (GE Helthcare, Little Chalfont, UK) and LAS1000 (FUJIFILM, Tokyo, Japan) were used. The images generated were subjected to densitometric analysis using ImageJ software (http://rsbweb.nih.gov/ij/index.html).

### Secondary metabolite extraction and High Resolution Mass Spectrometry (HRMS) analyses

For mass spectrometry analyses, extractions were performed according to [53] with modifications. Briefly, 100 mg of the fungal liophilized supernatant was extracted with 1 mL of MeOH during 40 minutes in ultrasonic bath. The extracts were centrifuged at 13000 rpm for 1 minute and the supernatants were collected and dried under N_2_. Crude extracts were resuspended in 600 uL MeOH and filtered in 0.22 µm. All the extractions were performed in triplicate.

The LC-HRMS analyses were performed in a liquid chromatography-mass spectrometer LC Agilent 1200 coupled to an Agilent iFunnel 6550 Q-ToF LC-MS with an ESI source. All the operation and spectra analyses were conducted using Agilent Mass Hunter Workstation Software. The parameters of MS analyses were set as follows: ESI source in positive mode; nebulizing gas temperature at 290 °C; capillary voltage at +3000 V; nozzle voltage at 320 V; drying gas flow of 12 ml min^−1^; nebulization gas pressure of 45 psi; auxiliary gas temperature at 350 °C; auxiliary gas flow of 12 ml min^−1^ and mass range of *m/z* 100-1700. The chromatographyc separation was performed on a Thermo Scientific Accucore C18 column (2.6 µm, 2.1 mm x 100 mm) and 0.1% formic acid (A) and acetonitrile (B) were used as mobile phase. The eluent profile (A:B) was: 0-10 min, gradient from 95:5 to 2:98; 10-15 min and final isocratic elution with 2:98. The flow rate was set at 0.2 mL min^-1^ and 1 µL of sample was injected in each injection.

The low-resolution mass spectrometry analyses were performed in an Acquity ultra-performance LC system coupled to a Quattro Micro triple quadrupole mass spectrometer from Waters. The same chromatographic conditions were applied as before. The mass spectrometer parameters were as follows: capillary voltage, 3.0 kV; cone voltage, 20 V; extractor voltage, 3 V; source temperature, 150 °C; desolvation temperature, 350 °C; and desolvation gas flow, 600 L/h. Dry nitrogen was used as desolvation and nebulization gas and argon was used as collision gas. All the data were acquired and processed using MassLynx v 4.1 software (Waters).

### Cell wall polysaccharides extraction and sugar quantification

Fungal cell wall polysaccharides were extracted from 10 mg dry-frozen biomass as described previously [54]. One milliliter of extracted samples were concentrated 10 x by liophilization, and sugars subsequently analyzed by high-performance liquid chromatography (HPLC) using a YoungLin YL9100 series system (YoungLin, Anyang, Korea) equipped with a YL9170 series refractive index (RI) detector at 40°C. Samples were loaded in a REZEX ROA (Phenomenex, USA) column (300 × 7.8 mm) at 85°C and eluted with 0.05 M sulfuric acid at a flow rate of 1.5 ml/min.

### Transmission Electron Microscopy (TEM) analysis of cell wall

Strains were grown statically from 1 x 10^7^ conidia at 37 °C in MM for 24 h. Mycelia were harvested and immediately fixed in 0.1 M sodium phosphate buffer (pH 7.4) containing 2.5% (v/v) of glutaraldehyde and 2% (w/v) of paraformaldehyde for 24 h at 4 °C. Samples were encapsulated in agar (2% w/v) and subjected to fixation (1% OsO_4_), contrasting (1% uranyl acetate), ethanol dehydration, and a two-step infiltration process with Spurr resin (Electron Microscopy Sciences) of 16 h and 3 h at RT. Additional infiltration was provided under vacuum at RT before embedment in BEEM capsules (Electron Microscopy Sciences) and polymerization at 60 °C for 72 h. Semithin (0.5-µm) survey sections were stained with toluidine blue to identify the areas of best cell density. Ultrathin sections (60 nm) were prepared and stained again with uranyl acetate (1%) and lead citrate (2%). Transmission electron microscopy (TEM) images were obtained using a Philips CM-120 electron microscope at an acceleration voltage of 120 kV using a MegaView3 camera and iTEM 5.0 software (Olympus Soft Imaging Solutions GmbH). Cell wall thickness of 100 sections of different germlings were measured at 23,500 x magnification and images analyzed with the ImageJ software [55]. Statistical differences were evaluated by using one-way analysis of variance (ANOVA) and Tukey’s *post hoc* test.

### Preparation of BMDMs and cytokine quantification

BMDMs were prepared and isolated as described previously [34, 56]. Bone marrow cells were obtained from femur and tibia of adult (6 to 8 weeks of age) male C57BL/6 mice. The BMDMs were plated on 48-well plates (1.5 x 10^6^ cells/ml; 7.5 x 10^5^ cells/well), and the cultures were maintained for 48 h for quantification of cytokines. The cytokine measurement was quantified using an enzyme-linked immunosorbent assay (ELISA) kit, according to the protocol of the manufacturer (BD Biosciences, Pharmigen, San Diego, CA, USA).

### Phagocytosis and killing assay

BMDMs were used in all phagocytosis and killing assays, as described previously [34, 56]. The phagocytic index was obtained using 2 x 10^5^ cells/well, plated on glass 13-mm-diameter coverslips placed on 24-well plates. BMDMs were treated with the conidia of fungi (2 x 10^5^ conidia; macrophages/conidia = 1:1) for 4 h at 37 °C with 5% CO_2_. An average of 100 macrophages was enumerated to determine the percentage of conidia that were ingested per macrophage. For the killing assay, the BMDMs were treated with 5 x 10^5^ conidia (macrophages/conidia = 1:1) for 48 h at 37 °C with 5% CO_2_. Serial dilutions were prepared using the lysate, and the cells were then plated, and incubated at 37 °C for 48 h. The viable fungi were enumerated, and the CFU counts per milliliter were calculated. The experiments were repeated three times, each performed in triplicate.

### Virulence analysis in *Galleria mellonella* models

The *Galleria mellonella* larvae were obtained by breeding adult larvae [57] weighing 275-330 mg, kept starving in Petri dishes at 37 ° in the dark for 24 h prior to infection. All selected larvae were in the final stage of larval (sixth) stage development. Fresh conidia of each strain of *A. fumigatus* were counted using a hemocytometer and the initial concentration of the conidia suspensions for the infections were 2 × 10^8^ conidia/ml. A total of 5 μl (1 × 10^6^ conidia/larvae) of each suspension was inoculated per larva. The control group was composed of larvae inoculated with 5 μl of PBS to observe death by physical trauma. The inoculum was performed using the Hamilton syringe (7000.5KH) through the last left prolegate. After infection, the larvae were kept at 37 °C in Petri dishes in the dark and scored daily. Larvae were considered dead due to lack of movement in response to touch. The viability of the inoculum administered was determined by plating a serial dilution of the conidia on YAG medium and incubating the plates at 37 °C for 72 h.

## Supporting Information

**Supplementary Table S1 -** Strains and plasmids used in this work.

**Supplementary Table S2 –** Values (nm) of cell wall thickness measured in hyphae (n = 100) and conidia (n= 100) by transmission electron microscopy.

**Supplementary Table S3 –** Identities (ID), Log2-fold change (FC), p-values and adjusted p-values (padj) of genes identified in the RNA-sequencing comparing the Δ*gprM* strain to the wild-type strain when grown for 24 h at 37 °C. The first tab shows the entire genome whereas the second and third tabs show up- and down-regulated genes respectively, with a -1 < log2FC < 1.

**Supplementary Table S4 –** Identities (ID), Log2-fold change (FC), p-values and adjusted p-values (padj) of genes identified in the RNA-sequencing comparing the Δ*gprJ* strain to the wild-type strain when grown for 24 h at 37 °C. The first tab shows the entire genome whereas the second and third tabs show up- and down-regulated genes respectively, with a -1 < log2FC < 1.

**Supplementary Table S5 –** Identities (ID), Log2-fold change (FC), p-values and adjusted p-values (padj) of genes identified in the RNA-sequencing comparing the *xylP::gprM* strain to the wild-type strain when grown for 4 h in xlose minimal medium after transfer from glucose minimal medium at 37 °C. The first tab shows the entire genome whereas the second and third tabs show up- and down-regulated genes respectively, with a -1 < log2FC < 1.

**Supplementary Table S6 –** Identities (ID), Log2-fold change (FC), p-values and adjusted p-values (padj) of genes identified in the RNA-sequencing comparing the *xylP::gprJ* strain to the wild-type strain when grown for 1 h in xlose minimal medium after transfer from glucose minimal medium at 37 °C. The first tab shows the entire genome whereas the second and third tabs show up- and down-regulated genes respectively, with a -1 < log2FC < 1.

**Supplementary Table S7 –** Identities (ID), Log2-fold change (FC), p-values and adjusted p-values (padj) of genes identified in the RNA-sequencing comparing the Δ*mpkB* strain to the wild-type strain when grown for 24 h at 37 °C. The first tab shows the entire genome whereas the second and third tabs show up- and down-regulated genes respectively, with a -1 < log2FC < 1.

**Supplementary Table S8 –** Species and class names for all fungi used in phylogenetic analysis of AsgA homologues.

**Supplementary Table S9 –** Primers used in this work.

**Supplementary Figure S1 –** The GPCR GprJ is involved in DHN-melanin production. The *A. fumigatus* Δ*gprJ* produces a dark pigment in the culture medium after growth in glucose minimal medium for 5 days at 37 °C. Deletion of the DHN-melanin polyketide synthase PksP and the GPCR GprM suppress the colouring of the medium as seen in the wild-type strain.

**Supplementary Figure S2 –** The GPCRs GprM and GprJ are not involved in glucose sensing in *A. fumigatus*. A. Dry weight of fungal biomass when strains were grown for 24 h and 48 h in glucose minimal medium (GMM). B. Glucose concentrations in culture supernatants, measured over a time period of 44 h, when strains were grown from spores in GMM. Standard deviations are shown for the average of three biological replicates.

**Supplementary Figure S3 –** Confirmation of strains by PCR (A, C, E) and Southern blots (B, D). A. Genomic DNA (gDNA) from the *A. fumigatus* wild type, Δ*gprJ* and Δ*gprJ::gprJ* strains were used for PCR amplification using the *gprJ 5’ Fw pRS426* and *gprJ 3’ Rv ext* (P7/P29) primers. DNA bands of 3.5kb (wild-type), 4.3kb (Δ*gprJ*) and 5.4kb (Δ*gprJ::gprJ* strain) were amplified. B. The gDNA from *A. fumigatus* wild type, *ΔgprJ* and *ΔgprJ::gprJ strains* was digested with the restriction enzyme *EcoRI* and a 1kb DNA fragment from the 5‘ noncoding region was used as a hybridization probe. Restriction digestion and hybridisation resulted in the detection of a 8.3 kb DNA band in the wild type strain, a 3.8 kb band in the Δ*gprJ* deletion strain and a 6.8 kb band in the Δ*gprJ::gprJ^+^* null mutant. C. gDNA from the *A. fumigatus* wild type and Δ*gprJ* Δ*gprM* strains was used for PCR amplification using the *gprJ 5’ Fw pRS426* and *gprJ 3’ Rv ext* (P7/P29) primers and the primers for confirming *gprJ* deletion (as described above). Deletion of *gprM* was confirmed using the *gprM 5′Fw ext* and *gprM 3′Rv pRS426* (P28/P16) primers. Fragments of 3.5 kb (using primers for *gprJ*) and 4.2 kb (using primers for *gprM*) were generated for the wild-type strain. In the double deletion strain, a 4.3 kb band (using primers for *gprJ* deletion) and a band of 4.4 kb (using primers for *gprM* deletion) were amplified. (D) The gDNA from the *A. fumigatus* wild type and the double deletion strains was digested with the restriction enzyme *Eco*RI and a 1kb DNA fragment from the 5‘ noncoding region of the *gprJ* gene was used as a hybridization probe. Restriction digestion and hybridisation resulted in the detection of a 8.3 kb fragment in the wild type strain and a 3.8 kb band in the *ΔgprJ ΔgprM* strain. In addition, the gDNA from the wild type and double deletion strains was digested with the restriction enzyme *Bam*HI and a 1kb DNA fragment from the 5‘ noncoding region of the *gprM* gene was used as a hybridization probe. A fragment of 4.6 kb and a band of 4.0 kb were detected in the wild type and *ΔgprJ ΔgprM* strains respectively. E. gDNA from the *A. fumigatus* wild type and *xylp::gprJ* strains were used for PCR amplification using the *ORF gprJ Fw xylp* and *gprJ 3’ Rv ext* (P21/P29) primers. A fragment of 2.5 kb was ampliefied in the wild type strain, wherease a 5.2 kb band was amplified in the positive *xylP::gprJ* strain.

## References

1. Bassetti M, Giacobbe DR, Grecchi C, Rebuffi C, Zuccaro V, Scudeller L. 2020. Performance of existing definitions and tests for the diagnosis of invasive aspergillosis in critically ill, adult patients: a systematic review with qualitative evidence synthesis. J Infect 81: 131–146. doi: 10.1016/j.jinf.2020.03.065.

2. Kosmidis C, Denning DW. 2015. The clinical spectrum of pulmonary aspergillosis. Postgrad Med J 91: 403–10. 10.1136/postgradmedj-2014-206291rep.

3. Brown GD, Denning DW, Gow NA, Levitz SM, Netea MG, White TC. 2012. Hidden killers: human fungal infections. Sci Transl Med 4: 165rv13. doi: 10.1126/scitranslmed.3004404.

4. Xue C, Hsueh YP, Heitman J. 2008. Magnificent seven: roles of G protein-coupled receptors in extracellular sensing in fungi. FEMS Microbiol Rev 32: 1010–1032. doi:10.1111/j.1574-6976.2008.00131.x

5. Kochman K. 2014. Superfamily of G-protein coupled receptors (GPCRs) – extraordinary and outstanding success of evolution. Postepy Higieny I Medycyny Doswiadczalnej 68: 1225–1237. doi: 10.5604/17322693.1127326.

6. Van Dijck P. 2009. Nutrient sensing G protein-coupled receptors: interesting targets for antifungals? Med Mycol 47: 671–680 (2009). doi: 10.3109/13693780802713349.

7. Brown NA, Schrevens S, van Dijck P, Goldman GH. 2018. Fungal G-protein-coupled receptors: mediators of pathogenesis and targets for disease control. Nat Microbiol 3: 402–414. doi:10.1038/s41564-018-0127-5.

8. Grice CM, Bertuzzi M, Bignell EM. 2013. Receptor-mediated signaling in *Aspergillus fumigatus*. Front Microbiol 4: 26. doi: 10.3389/fmicb.2013.00026.

9. Lafon A, Han KH, Seo JA, Yu JH, d’Enfert, C. 2006. G-protein and cAMP-mediated signal in *Aspergilli*: a genomic perspective. Fungal Genet Biol 43: 490–502. doi: 10.1016/j.fgb.2006.02.001.

10. Gehrke A, Heinekamp T, Jacobsen ID, Brakhage AA. 2010. Heptahelical receptors GprC and GprD of *Aspergillus fumigatus* are essential regulators of colony growth, hyphal morphogenesis, and virulence. Appl Environ Microbiol 76: 3989–3998. doi: 10.1128/AEM.00052-10.

11. Jung MG, Kim SS, Yu JH, Shin KS. 2016. Characterization of *gprK* encoding a putative hybrid G-protein-coupled receptor in *Aspergillus fumigatus*. PLoS One 11: e0161312. doi: 10.1371/journal.pone.0161312.

12. Manfiolli AO, Siqueira FS, Dos Reis TF, Van Dijck P, Schrevens S, Hoefgen S, Föge M, Straßburger M, de Assis LJ, Heinekamp T, Rocha MC, Janevska S, Brakhage AA, Malavazi I, Goldman GH, Valiante V. 2019. Mitogen-Activated Protein Kinase Cross-Talk Interaction Modulates the Production of Melanins in *Aspergillus fumigatus*. mBio 10: e00215–19. doi: 10.1128/mBio.00215-19. PMID: 30914505.

13. Heinekamp T, Thywißen A, Macheleidt J, Keller S, Valiante V, Brakhage AA. 2012. *Aspergillus fumigatus* melanins: interference with the host endocytosis pathway and impact on virulence. Front Microbiol 3:440. doi: 10.3389/fmicb.2012.00440.

14. Nosanchuk JD, Stark RE, Casadevall A. 2015. Fungal melanin: what do we know about structure? Front Microbiol 6:1463. doi: 10.3389/fmicb.2015.01463.

15. Zadra I, Abt B, Parson W, Haas H. 2000. *xylP* promoter-based expression system and its use for antisense downregulation of the *Penicillium chrysogenum* nitrogen regulator NRE. Appl Environ Microbiol 66: 4810–4816. doi: 10.1128/aem.66.11.4810-4816.2000.

16. Valiante V, Macheleidt J, Foge M, Brakhage AA. 2015. The *Aspergillus fumigatus* cell wall integrity signaling pathway: drug target, compensatory pathways, and virulence. Front Microbiol 6: 325. doi: 10.3389/fmicb.2015.00325.

17. Dos Reis TF, Mellado L, Lohmar JM, Silva LP, Zhou JJ, Calvo AM, Goldman GH, Brown NA. 2019. GPCR-mediated glucose sensing system regulates light-dependent fungal development and mycotoxin production. PLoS Genet 15: e1008419. doi: 10.1371/journal.pgen.1008419.

18. de Assis LJ, Manfiolli A, Mattos E, Fabri JHTM, Malavazi I, Jacobsen ID, Brock M, Cramer RA, Thammahong A, Hagiwara D, Ries LNA, Goldman GH. 2018. Protein kinase A and high-osmolarity glycerol response pathways cooperatively control cell wall carbohydrate mobilization in *Aspergillus fumigatus*. mBio 9: e01952–18. doi: 10.1128/mBio.01952-18.

19. Akache B, Turcotte B. 2002. New regulators of drug sensitivity in the family of yeast zinc cluster proteins. J Biol Chem 277: 21254–21260. doi:10.1074/jbc.M202566200.

20. van de Veerdonk FL, Gresnigt MS, Romani L, Netea MG and Latgé JP. 2017. *Aspergillus fumigatus* morphology and dynamic host interactions. Nat Rev Microbiol 15: 661–674. doi: 10.1038/nrmicro.2017.90.

21. Balloy V and Chignard M. 2009. The innate immune response to *Aspergillus fumigatus*. Microbes Infect, 11 (12): 919–27. doi: 10.1016/j.micinf.2009.07.002.

22. Brown NA and Goldman GH. 2016. The contribution of *Aspergillus fumigatus* stress responses to virulence and antifungal resistance. J. Microbiol 54: 243–253. doi: 10.1007/s12275-016-5510-4.

23. Affeldt KJ, Carrig J, Amare M, Keller NP. 2014. Global survey of canonical *Aspergillus flavus* G protein-coupled receptors. mBio 5: e01501–e1514. doi: 10.1128/mBio.01501-14.

24. Jiang C, Cao S, Wang Z, Xu H, Liang J, Liu H, Wang G, Ding M, Wang Q, Gong C, Feng C, Hao C, Xu JR. 2019. An expanded subfamily of G-protein-coupled receptor genes in *Fusarium graminearum* required for wheat infection. Nat Microbiol 4: 1582–1591. doi: 10.1038/s41564-019-0468-8.

25. Dilks T, Halsey K, De Vos RP, Hammond-Kosack KE, Brown NA. 2019. Non-canonical fungal G-protein coupled receptors promote *Fusarium* head blight on wheat. PLoS Pathog 15: e1007666. doi: 10.1371/journal.ppat.1007666.

26. Frawley D, Stroe MC, Oakley BR, Heinekamp T, Straßburger M, Fleming AB, Brakhage AA, Bayram Ö. 2020. The Pheromone Module SteC-MkkB-MpkB-SteD-HamE Regulates Development, Stress Responses and Secondary Metabolism in *Aspergillus fumigatus*. Front Microbiol 11:811. doi: 10.3389/fmicb.2020.00811.

27. Liebmann B, Gattung S, Jahn B, Brakhage AA. 2003. cAMP signaling in *Aspergillus fumigatus* is involved in the regulation of the virulence gene *pksP* and in defense against killing by macrophages. Mol Genet Genom 269: 420–35. doi: 10.1007/s00438-003-0852-0.

28. Grosse C, Heinekamp T, Kniemeyer O, Gehrke A, Brakhage AA. 2008. Protein kinase A regulates growth, sporulation, and pigment formation in *Aspergillus fumigatus*. Appl Environ Microbiol 74: 4923–4933. doi: 10.1128/AEM.00470-08.

29. Jain R, Valiante V, Remme N, Docimo T, Heinekamp T, Hertweck C, Gershenzon J, Haas H, Brakhage AA. 2011. The MAP kinase MpkA controls cell wall integrity, oxidative stress response, gliotoxin production and iron adaptation in *Aspergillus fumigatus*. Mol Microbiol 82: 39–53. doi: 10.1111/j.1365-2958.2011.07778.x.

30. Müller S, Baldin C, Groth M, Guthke R, Kniemeyer O, Brakhage AA, Valiante V. 2012. Comparison of transcriptome technologies in the pathogenic fungus *Aspergillus fumigatus* reveals novel insights into the genome and MpkA dependent gene expression. BMC Genomics 13: 519. doi: 10.1186/1471-2164-13-519.

31. Valiante V, Jain R, Heinekamp T, Brakhage AA. 2009. The MpkA MAP kinase module regulates cell wall integrity signaling and pyomelanin formation in *Aspergillus fumigatus*. Fungal Genet Biol 46: 909–918. doi: 10.1016/j.fgb.2009.08.005.

32. Rocha MC, Fabri JH, Franco de Godoy K, Alves de Castro P, Hori JI, Ferreira da Cunha A, Arentshorst M, Ram AF, van den Hondel CA, Goldman GH, Malavazi I. 2016. *Aspergillus fumigatus* MADS-Box Transcription Factor *rlmA* Is Required for Regulation of the Cell Wall Integrity and Virulence. G3 (Bethesda) 6: 2983–3002. doi: 10.1534/g3.116.031112.

33. Ries LNA, Rocha MC, de Castro PA, Silva-Rocha R, Silva RN, Freitas FZ, de Assis LJ, Bertolini MC, Malavazi I, Goldman GH. 2017. The *Aspergillus fumigatus* CrzA Transcription Factor Activates Chitin Synthase Gene Expression during the Caspofungin Paradoxical Effect. mBio 8: e00705–17. doi: 10.1128/mBio.00705-17.

34. de Castro PA, Colabardini AC, Manfiolli AO, Chiaratto J, Silva LP, Mattos EC, Palmisano G, Almeida F, Persinoti GF, Ries LNA, Mellado L, Rocha MC, Bromley M, Silva RN, de Souza GS, Loures FV, Malavazi I, Brown NA, Goldman GH. 2019. *Aspergillus fumigatus* calcium-responsive transcription factors regulate cell wall architecture promoting stress tolerance, virulence and caspofungin resistance. PLoS Genet 15: e1008551. doi: 10.1371/journal.pgen.1008551.

35. Mattos EC, Silva LP, Valero C, de Castro PA, Dos Reis TF, Ribeiro LFC, Marten MR, Silva-Rocha R, Westmann C, da Silva CHTP, Taft CA, Al-Furaiji N, Bromley M, Mortensen UH, Benz JP, Brown NA, Goldman GH. 2020. The *Aspergillus fumigatus* Phosphoproteome Reveals Roles of High-Osmolarity Glycerol Mitogen-Activated Protein Kinases in Promoting Cell Wall Damage and Caspofungin Tolerance. mBio 11(1): e02962–19. doi: 10.1128/mBio.02962-19.

36. Camacho C, Coulouris G, Avagyan V, Ma N, Papadopoulos J, Bealer K, Madden TL. 2009. BLAST+: architecture and applications. BMC Bioinformatics 10: 421. doi: 10.1186/1471-2105-10-421.

37. Steenwyk JL, Shen XX, Lind AL, Goldman GH, Rokas A. 2019. A Robust Phylogenomic Time Tree for Biotechnologically and Medically Important Fungi in the Genera *Aspergillus* and *Penicillium* mBio 10: e00925–19; doi: 10.1128/mBio.00925-19.

38. Katoh K, Standley DM. 2013. MAFFT multiple sequence alignment software version 7: improvements in performance and usability. Mol Biol Evol 30: 772–780. doi:10.1093/molbev/mst010.

39. Cock PJA, Antao T, Chang JT, Chapman BA, Cox CJ, Dalke A, Friedberg I, Hamelryck T, Kauff F, Wilczynski B, de Hoon MJL, 2009. Biopython: freely available Python tools for computational molecular biology and bioinformatics. Bioinformatics 25: 1422–1423. https://doi.org/10.1093/bioinformatics/btp163.

40. Capella-Gutiérrez S, Silla-Martínez JM, Gabaldón T. 2009. trimAl: a tool for automated alignment trimming in large-scale phylogenetic analyses. Bioinformatics 25: 1972–1973. doi:10.1093/bioinformatics/btp348.

41. Nguyen LT, Schmidt HA, von Haeseler A, Minh BQ. IQ-TREE: A Fast and Effective Stochastic Algorithm for Estimating Maximum-Likelihood Phylogenies. Molecular Biology and Evolution 32: 268–274. https://doi.org/10.1093/molbev/msu300.

42. Hoang DT, Vinh LS, Flouri T, Stamatakis A, von Haeseler A, Minh BQ. 2018. MPBoot: fast phylogenetic maximum parsimony tree inference and bootstrap approximation. BMC Evol Biol 18: 11. doi: 10.1186/s12862-018-1131-3.

43. Käfer E. 1977. Meiotic and mitotic recombination in *Aspergillus* and its chromosomal aberrations. Adv Genet 19: 33–131. doi: 10.1016/s0065-2660(08)60245-x.

44. Kubodera T, Yamashita N, Nishimura A. 2000. Pyrithiamine Resistance Gene (*ptrA*) of *Aspergillus oryzae*: Cloning, Characterization and Application as a Dominant Selectable Marker for Transformation. Bioscience, Biotechnology, and Biochemistry 64: 1416–1421. doi:10.1271/bbb.64.1416.

45. Colot HV, Park G, Turner GE, Ringelberg C, Crew CM, Litvinkova L, Weiss RL, Borkovich KA, Dunlap JC. 2006. A high-throughput gene knockout procedure for *Neurospora* reveals functions for multiple transcription factors. Proc Natl Acad Sci USA 103:10352–10357. doi: 10.1073/pnas.0601456103.

46. da Silva Ferreira ME, Kress MR, Savoldi M, Goldman MH, Härtl A, Heinekamp T, Brakhage AA, Goldman GH. 2006. The akuB(KU80) mutant deficient for nonhomologous end joining is a powerful tool for analyzing pathogenicity in *Aspergillus fumigatus*. Eukaryot Cell 5: 207–211. doi: 10.1128/EC.5.1.207-211.2006.

47. Malavazi I, Goldman GH. 2012. Gene disruption in *Aspergillus fumigatus* using a PCR-based strategy and in vivo recombination in yeast. Methods Mol Biol 845:99–118. doi: 10.1007/978-1-61779-539-8_7.

48. Furukawa T, van Rhijn N, Fraczek M, Gsaller F, Davies E, Carr P, Gago S, Fortune-Grant R, Rahman S, Gilsenan JM, Houlder E, Kowalski CH, Raj S, Paul S, Cook P, Parker JE, Kelly S, Cramer RA, Latgé JP, Moye-Rowley S, Bignell E, Bowyer P, Bromley MJ. 2020. The negative cofactor 2 complex is a key regulator of drug resistance in *Aspergillus fumigatus*. Nat Commun 11: 427. doi: 10.1038/s41467-019-14191-1.

49. Bolger AM, Lohse M, Usadel B. 2014. Trimmomatic: a flexible trimmer for Illumina sequence data. Bioinformatics 30: 2114–2120. doi: 10.1093/bioinformatics/btu170.

50. Kopylova E, Noe L, Touzet H. 2012. SortMeRNA: fast and accurate filtering of ribosomal RNAs in metatranscriptomic data. Bioinformatics 28: 3211–3217. doi: 10.1093/bioinformatics/bts611.

51. Liao Y, Smyth GK, Shi W. 2013. The Subread aligner: fast, accurate and scalable read mapping by seed-and-vote. Nucleic Acids Res 41: e108. doi: 10.1093/nar/gkt214.

52. Benjamini Y, Drai D, Elmer G, Kafkafi N, Golani I. 2001. Controlling the false discovery rate in behavior genetics research. Behav Brain Res 125: 279–84. doi: 10.1016/s0166-4328(01)00297-2.

53. Smedsgaard J. 1997. Micro-scale extraction procedure for standardized screening of fungal metabolite production in cultures. J Chromatogr A 760: 264–270. doi: 10.1016/s0021-9673(96)00803-5.

54. François JM. 2006. A simple method for quantitative determination of polysaccharides in fungal cell walls. Nat Protoc, 1: 2995–3000. doi: 10.1038/nprot.2006.457.

55. Schneider CA, Rasband WS and Eliceiri KW. 2012. NIH Image to ImageJ: 25 years of image analysis. Nat Methods 9: 671–5. doi: 10.1038/nmeth.2089.

56. Brauer VS, Pessoni AM, Bitencourt TA, de Paula RG, de Oliveira Rocha L, Goldman GH, Almeida F. 2020. Extracellular Vesicles from Aspergillus flavus Induce M1 Polarization *In Vitro*. mSphere 5: e00190–20. doi: 10.1128/mSphere.00190-20.

57. Fuchs BB, O’Brien E, Khoury JB, Mylonakis E. 2010. Methods for using *Galleria mellonella* as a model host to study fungal pathogenesis. Virulence 1: 475–82 doi:10.4161/viru.1.6.12985.

