## Supplementary Table S1 for "*Aspergillus fumigatus* G-protein coupled receptors GprM and GprJ are important for the regulation of the cell wall integrity pathway, secondary metabolite production, and virulence"

**Supplementary Table S1 - Strains and plasmids used in this work.**

| **Strain** | **Genotype** | **Reference** |
| --- | --- | --- |
| *A. fumigatus* |  |  |
| CEA17 | Δ*akuB pyrG^+^* | Da Silva Ferreira *et al.* (2006) |
| CEA17pyrG | Δ*akuB pyrG^-^* | Da Silva Ferreira *et al.* (2006) |
| Δ*gprM* | *ΔgprM::ptrA,* Δ*akuB* | Manfiolli *et al.* (2019) |
| Δ*gprM* | Δ*akuB* ^KU80^*;ΔgprM::prtA;* PT^R^ | Manfiolli et al., 2019 |
| Δ*gprM::gprM^+^* | Δ*akuB*^KU80^*;* Δ*gprM::prtA; gprM::gprM*-hph; Hyg^R^ | Manfiolli et al., 2019 |
| Δ*gprJ* | Δ*akuB*^KU80^ *ΔgprJ::prtA;* PT^R^ | This study |
| *ΔgprJ* pyrG^-^ | Δ*akuB*^KU80^ *gprJ::prtA;* pyrG-; PT^R^ | This study |
| Δ*gprJ::gprJ^+^* | Δ*akuB*^KU80^ *;*Δ*gprJ::prtA;pyrG-; gprJ::*pyrG; pyrG+ | This study |
| Δ*gprJ* Δ*gprM* | Δ*akuB*^KU80^ *;ΔgprJ::prtA; ΔgprM::pyrG;* PT^R^*,* pyrG+ | This study |
| *xylp::gprM* | Δ*akuB*^KU80^; *xylp*:: *gprM::*3HA::*prtA*; PT^R^ | Manfiolli et al., 2019 |
| *xylp::gprJ* | Δ*akuB*^KU80^ ;*xylp*::*gprJ*- 3HA::pyrG; pyrG+ | This study |
| Δ*gprJ* Δ*pksP* | Δ*gprJ pyrG^−^;* *pksP*::*pyrG*; PT^R^, PyrG+ | This study |
| Δ*mpkB* | *ΔmpkB::ptrA,* Δ*akuB* | Manfiolli *et al.* (2019) |
| Δ*gprA* | *ΔgprA::ptrA,* Δ*akuB* | Manfiolli *et al.* (2019) |
| Δ*gprB* | *ΔgprB::ptrA,* Δ*akuB* | Manfiolli *et al.* (2019) |
| Δ*gprC* | *ΔgprC::ptrA,* Δ*akuB* | Manfiolli *et al.* (2019) |
| Δ*gprD* | *ΔgprD::ptrA,* Δ*akuB* | Manfiolli *et al.* (2019) |
| Δ*gprH* | *ΔgprH::ptrA,* Δ*akuB* | Manfiolli *et al.* (2019) |
| Δ*gprI* | *ΔgprI::ptrA,* Δ*akuB* | Manfiolli *et al.* (2019) |
| Δ*gprJ* | *ΔgprJ::ptrA,* Δ*akuB* | Manfiolli *et al.* (2019) |
| Δ*gprK* | *ΔgprK::ptrA,* Δ*akuB* | Manfiolli *et al.* (2019) |
| Δ*gprM* | *ΔgprM::ptrA,* Δ*akuB* | Manfiolli *et al.* (2019) |
| Δ*gprO* | *ΔgprO::ptrA,* Δ*akuB* | Manfiolli *et al.* (2019) |
| Δ*gprP* | *ΔgprP::ptrA,* Δ*akuB* | Manfiolli *et al.* (2019) |
| Δ*nopA* | *ΔnopA::ptrA,* Δ*akuB* | Manfiolli *et al.* (2019) |
| *S. cerevisiae* |  |  |
| SC9721 | *MATa his3-D200 URA 3-52 leu2D1 lys2D202 trp1D63)* | FGSC |
| Plasmids |  |  |
| pCDA21 | *Zeo::pyr amp^R^* | Chaveroche et al., 2000 |
| pSK275 | Amp^R^, ptrA cassette | Pearson et al., 2001 |
| pRS426 | Amp^R^ *lacZ* URA3 | Teepe et al., 2007 |
| pYES-xyl^P^-hph-devR | Amp^R^ *lacZ* | Özgur Bayram lab |
| pOB430 | Amp^R^ *lacZ* | Özgur Bayram lab |

FGSC=Fungal Genetics Stock Center ([www.fgsc.net](http://www.fgsc.net)); Hyg^R^, hygromycin; PT^R^, pyrithiamine

**References**

Da Silva Ferreira ME, Kress MR, Savoldi M, Goldman MH, Härtl A, Heinekamp T, Brakhage AA, Goldman GH. 2006. The akuB(KU80) mutant deficient for nonhomologous end joining is a powerful tool for analysing pathogenicity in A*spergillus fumigatus*. Eukaryot Cell 5:207-211.

Manfiolli AO, Siqueira FS, dos Reis TF, Van Dijck P, Schrevens S, Hoefgen S, Föge M, Straßburger M, de Assis LJ, Heinekamp T, Rocha MC, Janevska S, Brakhage AA, Malavazi I, Goldman GH, Valiante V. 2019. Mitogenactivated protein kinase cross-talk interaction modulates the production of melanins in Aspergillus fumigatus. mBio 10:e00215-19. https://doi.org/10.1128/mBio.00215-19.

Pearson G, Robinson F, Beers Gibson T, Xu BE, Karandikar M, Berman K, Cobb MH. 2001. Mitogen-activated protein (MAP) kinase pathways: regulation and physiological functions. Endocr Rev 22:153–183. doi:10.1210/edrv.22.2.0428

Teepe, A. G., Loprete, D. M., He, Z., Hoggard ,T. A., Hill, T.W. The protein kinase C orthologue PkcA plays a role in cell wall integrity and polarized growth in *Aspergillus nidulans*. *Fungal Genet Biol*. 44, 554–562 (2007)

Chaveroche MK, Ghigo JM, d’Enfert C. A rapid method for efficient gene replacement in the filamentous fungus Aspergillus nidulans. Nucleic Acids Res. 2000;28:E97. doi: 10.1093/nar/28.22.e97.

d’Enfert C. 1996. Selection of multiple disruption events in *Aspergillus fumigatus* using the orotidine-5=-decarboxylase gene, pyrG, as a unique transformation marker. Curr Genet 30:76 – 82. https://doi.org/10.1007/ s002940050103.
