## Supplementary Table S9 for "*Aspergillus fumigatus* G-protein coupled receptors GprM and GprJ are important for the regulation of the cell wall integrity pathway, secondary metabolite production, and virulence"

**Supplementary Table S9. Primers used in this work**

|  | **Primer name** | **Primer sequence (5´- 3´)** |
| --- | --- | --- |
| P1 | gprJ 5´Fw | ATCAGGGAGAAGCTGGTGGCG |
| P2 | gprJ 5’ Rv ptrA | GGCCTGAGTGGCCATCGAATTCAGAGCCCAGAAGACCAGAGG |
| P3 | gprJ 3’ Fw ptrA | GAGGCCATCTAGGCCATCAAGCTTGGCTCTTTGGGCACGATGG |
| P4 | gprJ_3´rev | ATCCGTTCGACGTTTTGTTTCC |
| P5 | PtrA_fw_II | GAATTCGATGGCCACTCAGGCC |
| P6 | PtrA_Rv_II | GAGGCCATCTAGGCCATCAAGC |
| P7 | gprJ 5’ Fw pRS426 | GTAACGCCAGGGTTTTCCCAGTCACGACGATCAGGGAGAAGCTGGTGGCG |
| P8 | gprJ ORF Rv pyrG | CAGTGCCTCCTCTCAGACAGAATATCCTTATCGAAGAGTGAGATG |
| P9 | gprJ 3’ Fw pyrG | GAGCATTGTTTGAGGCGAATTCCATCTCACTCTTCGATAAGG |
| P10 | gprJ 3´Rv pRS426 | GCGGATAACAATTTCACACAGGAAACAGCCGCTTACGCTGTGACTTATTTC |
| P11 | pyrG F | ATTCTGTCTGAGAGGAGGCACTGATGCG |
| P12 | pyrG R | GAATTCGCCTCAAACAATGCTCTTCACC |
| P13 | gprM 5´F pRS426 | GTAACGCCAGGGTTTTCCCAGTCACGACGCCACGGACACATTTTTACG |
| P14 | gprM 5´R pyrG | CAGTGCCTCCTCTCAGACAGAATACTGCCGGTCTTGTTCGTGT |
| P15 | gprM 3´Fw pyrG | GAGCATTGTTTGAGGCGAATTCACTGGCGTCTTTTTTTTTGG |
| P16 | gprM 3´Rv pRS426 | GCGGTTAACAATTTCTCTCTGGAAACAGCTACTAGTCTTGCCCCGACTA |
| P17 | xylP Fw | GCACTGATGCGAGCAACAGTATG |
| P18 | xylP_Rv | CGACTCGAAGAACCAACC |
| P19 | HA-trpC-pyrG Fw | GGAGGTGGTAGCGGTGGT |
| P20 | HA-trpC-pyrG Rv | CTGTCTGAGAGGAGGCACTGAT |
| P21 | ORF gprJ Fw xyl^P^ | CGACTCGAAGAACCAACCATGTCCCTCCCATTGGTGCAGG |
| P22 | gprJ ORF Rv HA-trpC-pyrG | ACCACCGCTACCACCTCCCGACACTGCAGTCGTGAGCTG |
| P23 | xyl^P^ gprJ 5´Rv | CATACTGTTGCTCGCATCAGTGCACCCGATCCCTGCGATTACAC |
| P24 | pyrG gprJ 3´ Fw | ATCAGTGCCTCCTCTCAGACAGCATCTCACTCTTCGATAAGG |
| P25 | pRS gprJ 3´ Rv | GCGGATAACAATTTCACACAGGAAACAGCCGCTTACGCTGTGACTTATTTC |
| P26 | gprJ SYBR fw | GCGGATGGTGTGCTCATCGCGC |
| P27 | gprJ SYBR Rv | GCGAAGCCGGACCCCGTTCC |
| P28 | gprM 5´Fw ext | ACTGCATTGATCTCGGGC |
| P29 | gprJ 3’ Rv ext | CCCGACCATCTCCATCGACAAGAC |
| P30 | pksP 5’ Fw pRS426 | GTAACGCCAGGGTTTTCCCAGTCACGACGAACCTGAATCCTTGGCACTG |
| P31 | pksP 5’ Rv pyrG | CAGTGCCTCCTCTCAGACAGAATGGCGAGTGGTTTGCGCGGCC |
| P32 | pksP 3’ Fw pyrG | GAGCATTGTTTGAGGCGAATTCTGGGGTGAGTTCCTAGGTTT |
| P33 | pksP 3’ Rv pRS426 | GCGGTTAACAATTTCTCTCTGGAAACAGCTGTATTTTGTATGTTCAGCCTT |
| P34 | pksP 5’ Fw ext | CGCCCCGACCCTTACCGTGG |
| P35 | AFUB_019790 5’ Fw | CCACCTCTCGCCAAC |
| P36 | AFUB_019790 3’ Rv | GTTCCAGTAGCATAGGTCC |
| P37 | AFUB_019790 5’ Fw ext | GACCATCAGAAACTCGGAA |
