## Supplementary figures and images for "*Aspergillus fumigatus* G-protein coupled receptors GprM and GprJ are important for the regulation of the cell wall integrity pathway, secondary metabolite production, and virulence"

### Supplementary Figure S1

#### Slide 1
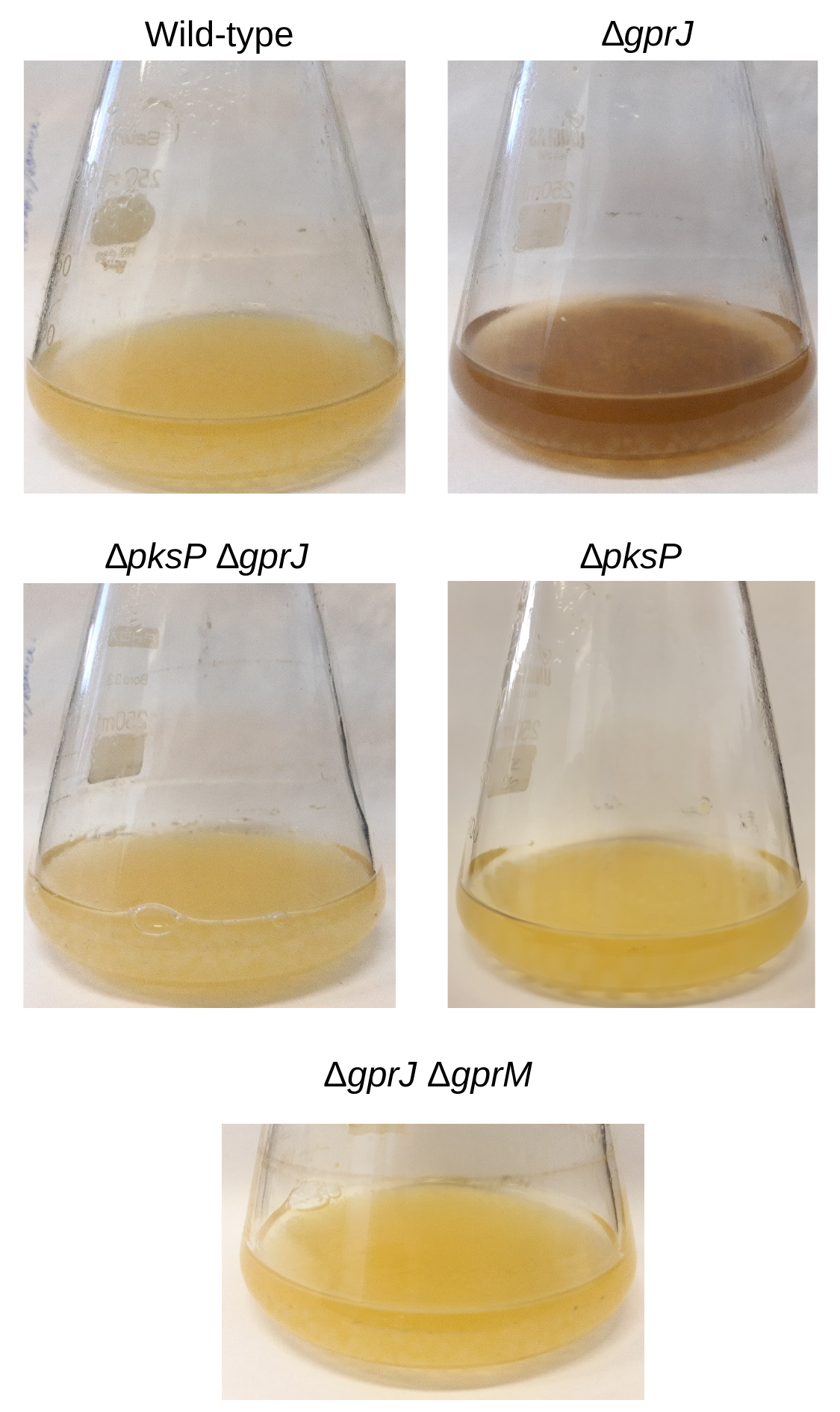

∆gprJ
Wild-type
∆pksP ∆gprJ
∆pksP
ΔgprJ ΔgprM
