## Supplementary Figure S2 for "*Aspergillus fumigatus* G-protein coupled receptors GprM and GprJ are important for the regulation of the cell wall integrity pathway, secondary metabolite production, and virulence"

### Slide 1
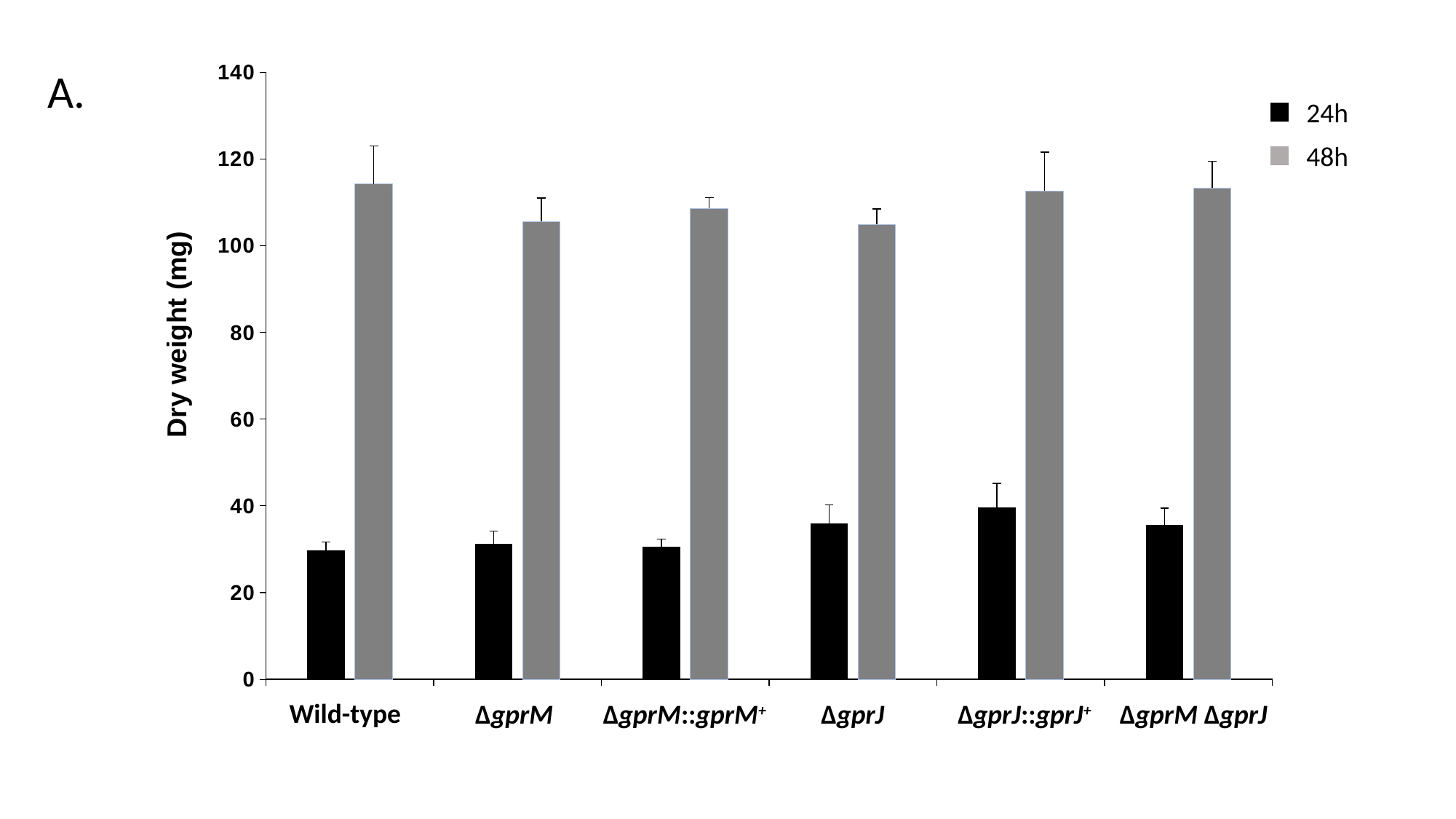

#### Chart
| Category | 24h | 48h |
|---|---|---|
| WT | 29.666666666666668 | 114.3333333333333 |
| DgprM | 31.333333333333318 | 105.66666666666667 |
| gprM+ | 30.666666666666668 | 108.66666666666667 |
| DgprJ | 36.0 | 105.0 |
| gprJ+ | 39.666666666666636 | 112.66666666666667 |
| DgprMDgprJ | 35.666666666666636 | 113.3333333333333 |A.
24h
48h
Dry weight (mg)
Wild-type
ΔgprM
ΔgprM::gprM^+^
ΔgprJ
ΔgprJ::gprJ^+^
ΔgprM ΔgprJ

### Slide 2
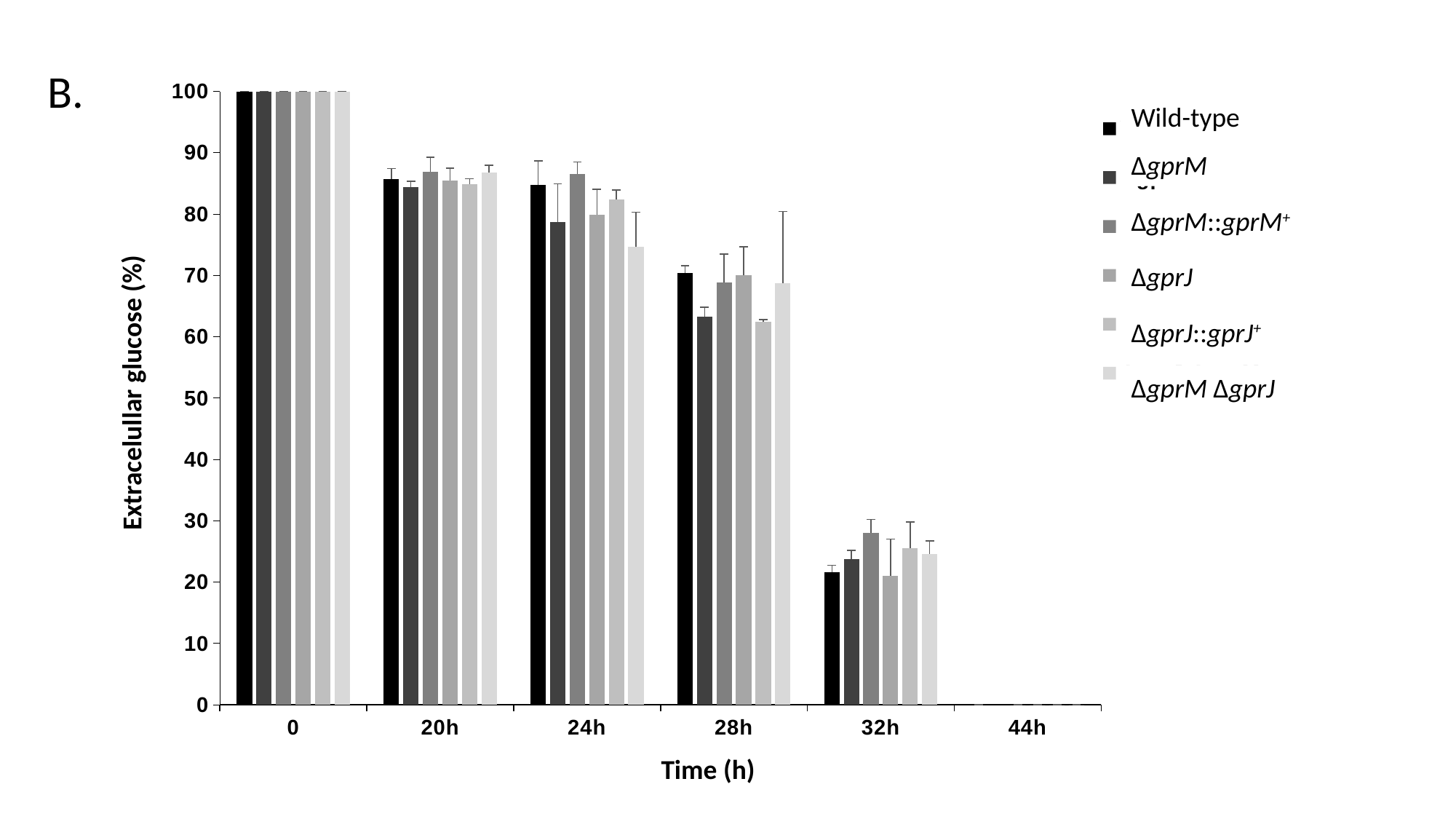

B.
[unsupported chart]
Wild-type
ΔgprM
ΔgprM::gprM^+^
ΔgprJ
ΔgprJ::gprJ^+^
ΔgprM ΔgprJ
Extracelullar glucose (%)
Time (h)
