## Supplementary Figure S3 for "*Aspergillus fumigatus* G-protein coupled receptors GprM and GprJ are important for the regulation of the cell wall integrity pathway, secondary metabolite production, and virulence"

### Slide 1
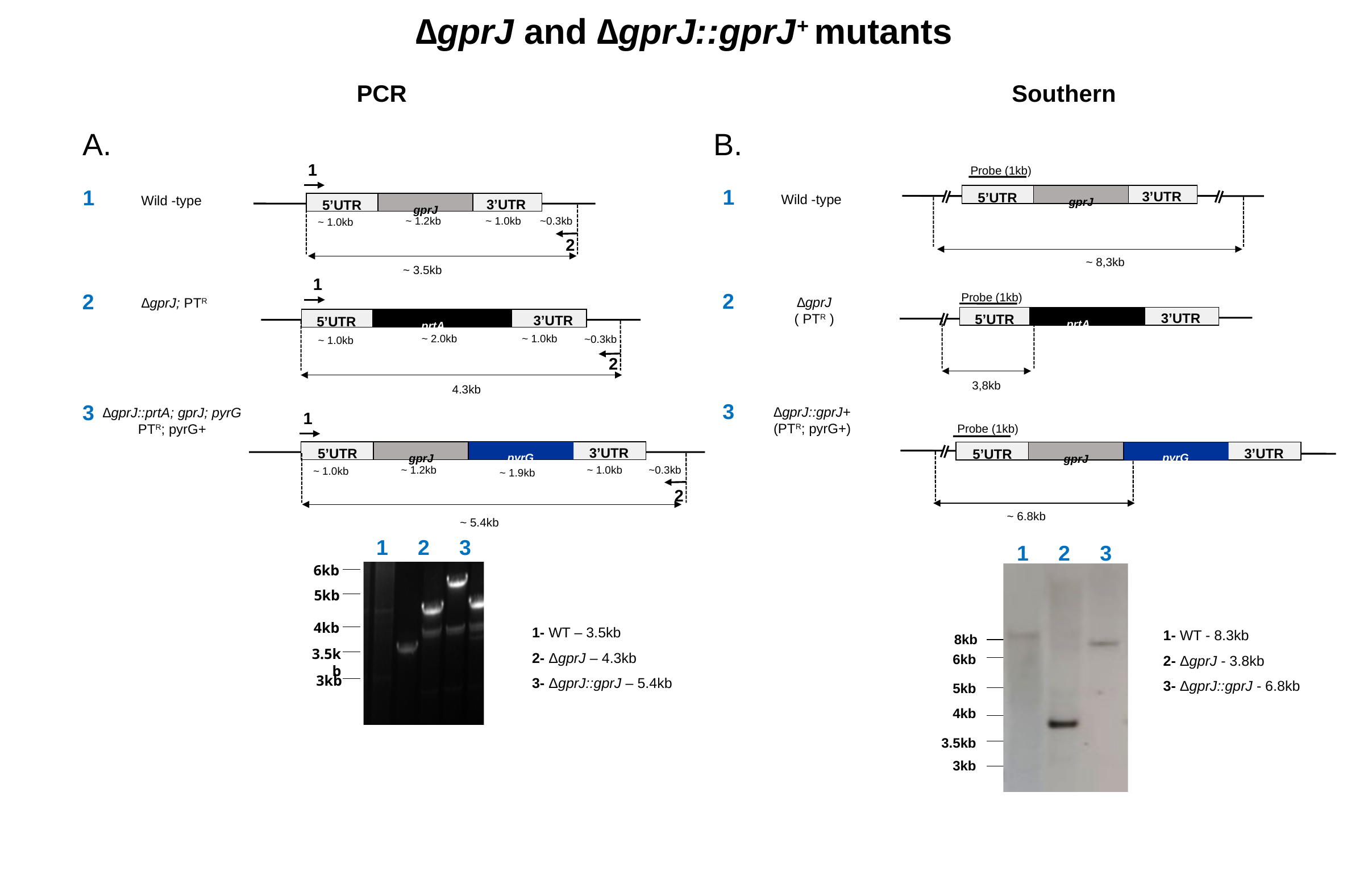

∆gprJ and ∆gprJ::gprJ^+^ mutants
PCR
Southern
A.
B.
1
3’UTR
5’UTR
^gprJ^
~ 3.5kb
Probe (1kb)
1
1
3’UTR
5’UTR
Wild -type
Wild -type
^gprJ^
~ 1.2kb
~ 1.0kb
~0.3kb
~ 1.0kb
2
~ 8,3kb
1
3’UTR
5’UTR
^prtA^
~ 2.0kb
~ 1.0kb
~0.3kb
~ 1.0kb
2
4.3kb
2
2
Probe (1kb)
3’UTR
5’UTR
^prtA^
3,8kb
∆gprJ
( PT^R^ )
∆gprJ; PT^R^
3
3
∆gprJ::gprJ+
(PT^R^; pyrG+)
∆gprJ::prtA; gprJ; pyrG
PT^R^; pyrG+
1
3’UTR
5’UTR
^pyrG^
^gprJ^
~ 1.2kb
~ 1.0kb
~0.3kb
~ 1.0kb
2
~ 5.4kb
Probe (1kb)
3’UTR
5’UTR
^pyrG^
^gprJ^
~ 6.8kb
~ 1.9kb
 1 2 3
1 2 3
6kb
5kb
1- WT – 3.5kb
2- ΔgprJ – 4.3kb
3- ΔgprJ::gprJ – 5.4kb
1- WT - 8.3kb
2- ΔgprJ - 3.8kb
3- ΔgprJ::gprJ - 6.8kb
4kb
8kb
3.5kb
6kb
3kb
5kb
4kb
3.5kb
3kb

### Slide 2
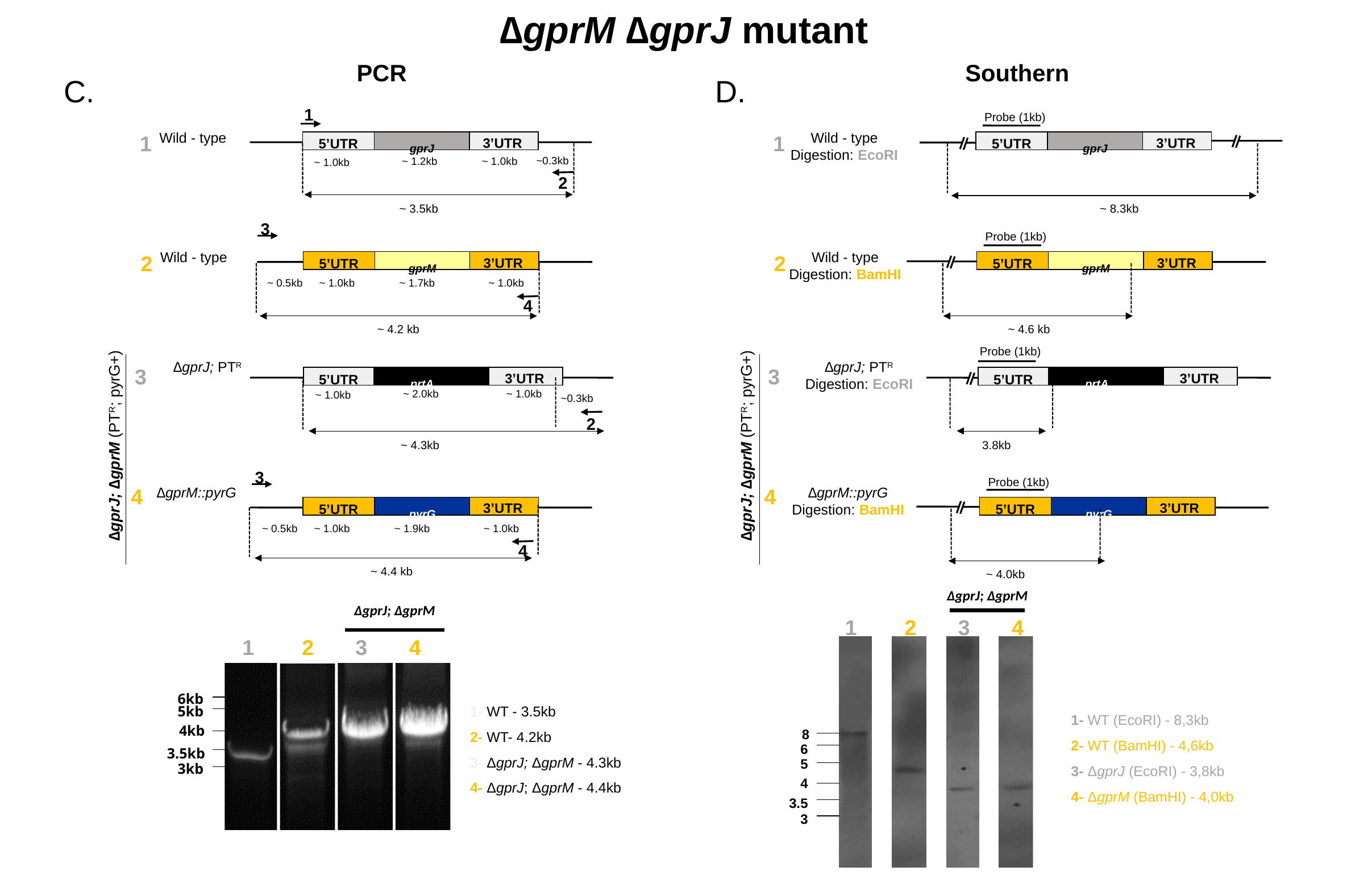

∆gprM ∆gprJ mutant
PCR
Southern
C.
D.
1
Probe (1kb)
1
Wild - type
Digestion: EcoRI
3’UTR
5’UTR
^gprJ^
~ 8.3kb
Wild - type
1
3’UTR
5’UTR
^gprJ^
~0.3kb
~ 1.2kb
~ 1.0kb
~ 1.0kb
2
~ 3.5kb
3
Probe (1kb)
2
Wild - type
Digestion: BamHI
3’UTR
5’UTR
^gprM^
~ 4.6 kb
Wild - type
2
3’UTR
5’UTR
^gprM^
~ 0.5kb
~ 1.0kb
~ 1.7kb
~ 1.0kb
4
~ 4.2 kb
Probe (1kb)
∆gprJ; PT^R^
Digestion: EcoRI
3’UTR
5’UTR
^prtA^
3.8kb
∆gprJ; PT^R^
3
3
3’UTR
5’UTR
^prtA^
~ 2.0kb
~ 1.0kb
~ 1.0kb
~0.3kb
2
∆gprJ; ∆gprM (PT^R^; pyrG+)
∆gprJ; ∆gprM (PT^R^; pyrG+)
~ 4.3kb
3
∆gprM::pyrG
∆gprM::pyrG
Digestion: BamHI
4
3’UTR
5’UTR
^pyrG^
~ 4.0kb
Probe (1kb)
4
3’UTR
5’UTR
^pyrG^
~ 0.5kb
~ 1.0kb
~ 1.9kb
~ 1.0kb
4
~ 4.4 kb
∆gprJ; ∆gprM
∆gprJ; ∆gprM
 1 2 3 4
8
6
5
4
3.5
3
 1 2 3 4
6kb
1- WT - 3.5kb
2- WT- 4.2kb
3- ΔgprJ; ΔgprM - 4.3kb
4- ΔgprJ; ΔgprM - 4.4kb
5kb
1- WT (EcoRI) - 8,3kb
2- WT (BamHI) - 4,6kb
3- ΔgprJ (EcoRI) - 3,8kb
4- ΔgprM (BamHI) - 4,0kb
4kb
3.5kb
3kb

### Slide 3
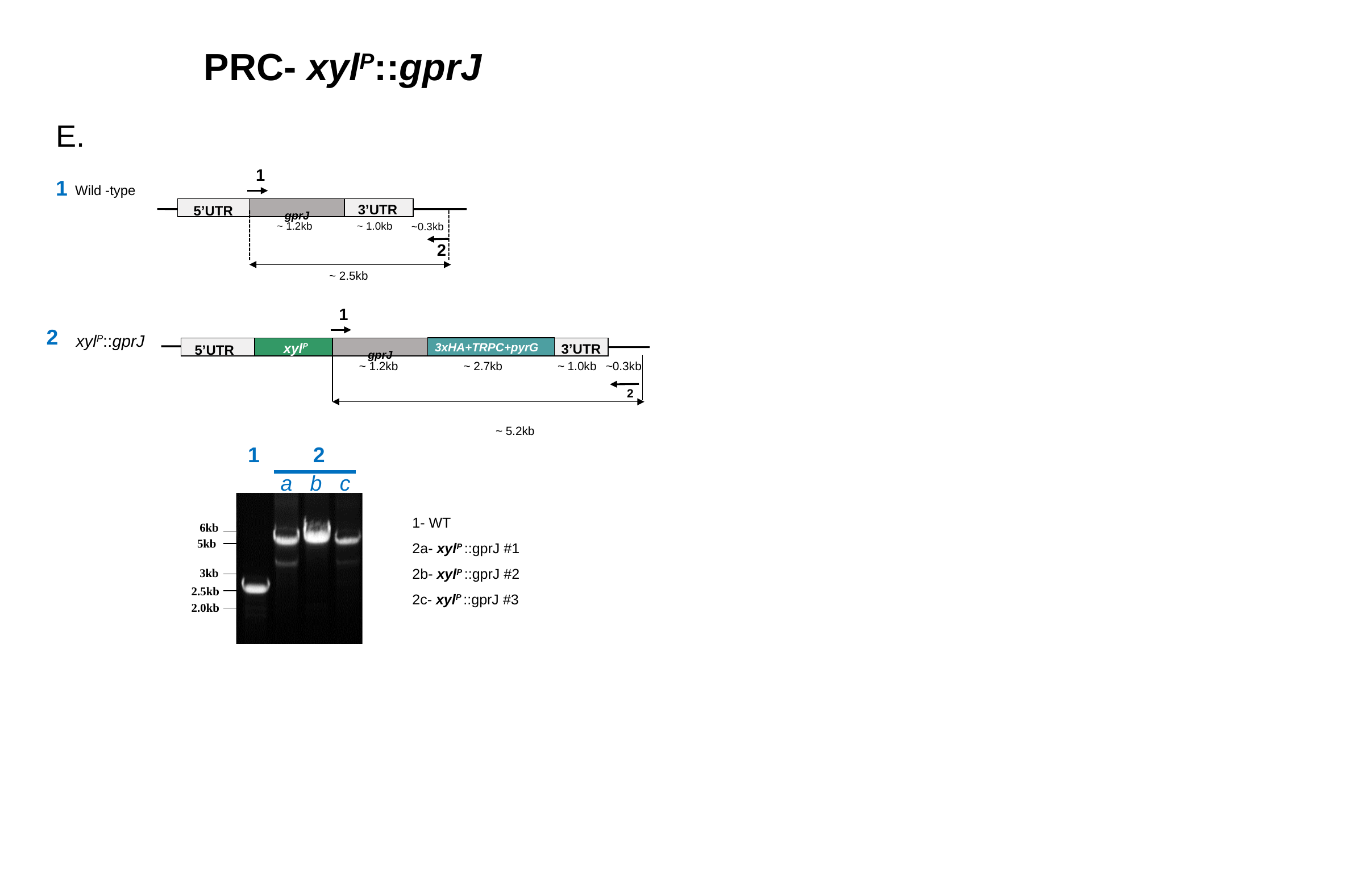

PRC- xyl^P^::gprJ
E.
1
3’UTR
5’UTR
^gprJ^
~ 2.5kb
1
Wild -type
~ 1.2kb
~ 1.0kb
~0.3kb
2
1
2
xyl^P^::gprJ
xyl^P^
3xHA+TRPC+pyrG
3’UTR
5’UTR
^gprJ^
~ 1.2kb
~ 2.7kb
~ 1.0kb
~0.3kb
2
~ 5.2kb
1 2
a b c
1- WT
2a- xyl^P^ ::gprJ #1
2b- xyl^P^ ::gprJ #2
2c- xyl^P^ ::gprJ #3
6kb
5kb
3kb
2.5kb
2.0kb
